# Integrative multi-omics analyses reveal multi-modal FOXG1 functions acting on epigenetic processes and in concert with NEUROD1 to regulate synaptogenesis in the mouse hippocampus

**DOI:** 10.1101/2021.10.25.465562

**Authors:** Ipek Akol, Stefanie Heidrich, Darren ÓhAilín, Christine Hacker, Alejandro Villarreal, Tudor Rauleac, Chiara Bella, Andre Fischer, Thomas Manke, Tanja Vogel

**Author notes:** Corresponding author: Institute for Anatomy and Cell Biology, Dept. Molecular Embryology, Medical Faculty, University of Freiburg, Albertstr. 17, 79104 Freiburg, Germany.

## Abstract

**Background:** FOXG1 has important functions for neuronal differentiation and balances excitatory/inhibitory network activity. Mutations in the human *FOXG1* gene cause a rare neurodevelopmental disorder, FOXG1-syndrome, which manifests differing phenotypes, including severe cognitive dysfunction, microencephaly, social withdrawal, and communication and memory deficits. Changes at the molecular level underlying these functional abnormalities upon *FOXG1* haploinsufficiency are largely unexplored, in human patients as well as in animals modelling the debilitating disease.

**Methods:** We present multi-omics data and explore comprehensively how FOXG1 impacts neuronal maturation at the chromatin level in the adult mouse hippocampus. We used RNA-, ATAC- and ChIP-sequencing of primary hippocampal neurons and co-immunoprecipitation to explore various levels of epigenetic changes and transcription factor networks acting to alter neuronal differentiation upon reduction of FOXG1.

**Results:** We provide the first comprehensive multi-omics data set exploring FOXG1 presence at the chromatin and identifying the consequences of reduced FOXG1 expression in primary hippocampal neurons. Analyzing the multi-omics data, our study reveals that FOXG1 uses various different ways to regulate transcription at the chromatin level. On a genome-wide level, FOXG1 (i) both represses and activates transcription, (ii) binds mainly to enhancer regions, and (iii) bidirectionally alters the epigenetic landscape in regard to levels of H3K27ac, H3K4me3, and chromatin accessibility. Genes affected by the chromatin alterations upon FOXG1 reduction impact synaptogenesis and axonogenesis. This finding emphasizes the importance of FOXG1 to integrate and coordinate transcription of genes necessary for proper neuronal function by acting on a genome-wide level. Interestingly, FOXG1 acts through histone deacetylases (HDACs) and inhibition of HDACs partly rescued transcriptional alterations observed upon FOXG1 reduction. On a more detailed level of analysis, we show that FOXG1 (iv) operates synergistically with NEUROD1. Interestingly, we could not detect a clear hierarchy of these two key transcription factors, but instead provide first evidence that they act in highly concerted and orchestrated manner to control neuronal differentiation.

**Conclusions:** This integrative and multi-omics view of changes upon FOXG1 reduction reveals an unprecedented multimodality of FOXG1 functions converging on neuronal maturation, fueling novel therapeutic options based on epigenetic drugs to alleviate, at least in part, neuronal dysfunctions.

## Background

Forkhead box G1 (FOXG1)-syndrome is a rare, congenital neurodevelopmental disorder (OMIM #613454), and patients present with a complex phenotypic spectrum encompassing microcephaly, seizures, scoliosis, abnormal movement, coordination and communication disorders, disrupted circadian rhythm, social withdrawal, avoidance of eye contact, indifference to visual/auditory stimuli and severe cognitive dysfunction (1–3). FOXG1-syndrome also falls under the umbrella of autism-spectrum disorders (4). As of yet, therapeutic options are limited for patients with FOXG1-syndrome. To overcome this limitation, research on FOXG1 in model organisms, such as the mouse, needs to unravel its seemingly diverse molecular functions (5).

In the mouse, FOXG1 acts early in central nervous system (CNS) development, where it is recognized as one key determinant in forebrain development (6). In the developing ventral and dorsal telencephalon, FOXG1 affects both progenitor proliferation and differentiation, and thus strongly impacts corticogenesis (6–11).

Despite a considerable body of data based on cortical development, less information is available describing FOXG1 functions in the postnatal brain or on the consequences of impaired FOXG1 expression for neuronal functions. In mature cerebellar neurons, FOXG1 interacts with one of two MECP2 isoforms (MECP2-e2) to prevent cell death (12). In the postnatal hippocampus, where FOXG1 is strongly expressed in the dentate gyrus (DG), FOXG1 also prevents cell death of postnatally born DG neurons (13). Reduced levels of FOXG1 in the mouse hippocampus impact the animal’s behavior, as *Foxg1*-haploinsufficient mice show hyperactivity, impaired habituation in open field tests, reduced performance in contextual fear conditioning (14) and autism-like features (15). Additionally, FOXG1 fosters hippocampal progenitor proliferation and differentiation (13, 14). Recent studies in mice show altered electrophysiological properties in neurons upon FOXG1 reduction or increase, disbalanced neuronal functions of excitation and inhibition (which is an important basis for impaired behavior), stunted dendritic complexity and reduced spine densities (15–19). While these studies demonstrate that features of FOXG1 alterations in mice largely reflect those seen in human FOXG1-patients, the functions of FOXG1 are yet to be fully elucidated. Indeed, several of the mouse studies relied on the complete loss of both *Foxg1* alleles. Thus, the data retrieved from these models cannot fully tell us whether loss of a single *Foxg1* allele affects, for example, neuronal differentiation. Instead, using either heterozygous animals or models with reduced levels of FOXG1 would advance our understanding of the alterations behind the human patient’s conditions. Furthermore, the molecular mechanisms of FOXG1-syndrome have been scantly characterized. Although FOXG1 has been recognized as a key transcription factor (TF) of forebrain development for many years, in-depth analyses of chromatin-related FOXG1 functions during forebrain development are still limited (6,8,20) and chromatin-independent functions of FOXG1 are just emerging (21, 22). This lack of a high-resolution view on how FOXG1 operates molecularly hampers comprehensive mechanistic insights into its functions.

Mouse mutants and human features of FOXG1-syndrome indicate that impaired FOXG1 function in the adult hippocampus might account for some of the phenotypes observed, as the hippocampus is a major center for learning and memory, and it is a hotspot for seizures caused by misbalanced neuronal activity (22). Thus, to resolve some of the open questions mentioned above, we studied (i) mouse hippocampal neurons with reduced FOXG1 levels, and used (ii) a multi-omics approach to unravel FOXG1 functions at the chromatin level. We focused particularly on the impact of FOXG1 on neuronal differentiation to establish mechanistic links to the observed and reported phenotypic alterations in FOXG1-mutant animals or humans. Our data show that FOXG1 mainly binds to enhancer regions and reconfigures the epigenetic landscape by altering chromatin accessibility, and increasing and decreasing H3K27ac and H3K4me3 marks at putative enhancers. We further show that FOXG1 cooperates with HDACs and NEUROD1 to increase and decrease expression of genes that affect terminal differentiation/maturation of neurons.

## Methods

### Mice

All mouse experiments were approved by the animal welfare committees of the respective authorities. *Foxg1*^cre/+^ mice were maintained in a C57Bl/6 background.

### Mouse hippocampus isolation and primary neuronal culture

Hippocampi of E18.5 C57Bl/6 embryos (Charles River) were dissected and collected in 10 ml Hanks’ Balanced Salt Solution (HBSS, Fisher Scientific) and dissociated in 0.25% Trypsin/EDTA (Fisher Scientific) solution at 37 °C for 10 min. Dissociation was stopped by adding 10% fetal bovine serum (FBS, Fisher Scientific). Cells were collected by centrifugation and cultured in Neurobasal (NB)-complete medium, which consists of neurobasal medium (Fisher Scientific) supplemented with B27 (Fisher Scientific), L-glutamine (0.5 mM, Fisher Scientific), penicillin-streptomycin-neomycin (1X PSN, Fisher Scientific), apo-transferrin (5 µg/ml, Sigma), superoxide-dismutase (0.8 µg/ml, Sigma) and glutathione (1 µg/ml, Sigma). Cells were seeded on poly-L-ornithine (0.1 mg/ml, Sigma) and laminin (1 µg/ml, Sigma) coated 24-well plates (Corning).

### Viral transduction and selection of primary neurons

Lentiviral particles were prepared using plko.1-CMV.Puro-tGFP-shNeurod1, plko.1-CMV.Puro-tGFP-shFoxg1 or plko.1-CMV.Puro-tGFP-shLuciferase (Genscript) plasmids, according to the protocol described previously (23, 24). Cells were transduced with lentiviral particles on day-in-vitro (DIV) 1. At DIV4, transduced cells were selected using 0.3 µg/ml puromycin (P9620, Sigma) and cell proliferation was inhibited by addition of 2 µM Arabinocytosine (AraC, C1768, Sigma), while performing a half-volume medium change. Medium was changed again at DIV7 and DIV9 including 0.3 µg/ml puromycin and 2 µM AraC.

### RNA isolation from tissue or primary hippocampal neurons

Hippocampi were frozen in liquid nitrogen immediately after dissection, and stored at −80°C until RNA extraction. Total RNA was isolated using RNeasy RNA isolation kit (Qiagen) according to the instructions of the manufacturer. An on-column DNAse digestion was routinely performed. Isolated RNA was kept at −80°C until following qRTPCR experiments.

Cells were harvested at DIV7 or DIV11 in buffer RLT (RNeasy RNA isolation kit, Qiagen) for total RNA extraction according to manufacturer’s instructions. Total RNA was kept at −80°C until subsequent reverse transcription.

### Reverse transcription and qRTPCR

1 µg of total RNA was reverse transcribed using RevertAid MMuLV reverse transcriptase kit (Fermentas, Thermo Scientific). qRTPCR analysis was performed on a CFX-Connect Real-Time PCR detection system (Bio-Rad) using GoTaq qPCR Master Mix (Promega). Primers used had an efficiency level between 85% and 110%. qRTPCR results were analyzed using the ΔΔCt method with GAPDH as internal standard. GraphPad Prism software was used for plotting the bar graphs and statistical analyses. Values in bar graphs are expressed as average ± SEM.

### RNA-seq

Total RNA extracted from mouse hippocampi or DIV7 primary hippocampal neurons were used for RNA-seq. Libraries were prepared using the TruSeq total RNA sample preparation kit from Illumina, following the manufacturer’s instructions. The procedure included depletion of rRNA prior double-stranded cDNA synthesis and library preparation. Samples were sequenced on Illumina HiSeq2500 as paired-end 100 bp reads.

### ChIP-seq

#### In vivo hippocampal tissue samples

CA and DG regions of the hippocampus were dissected separately and snap frozen in liquid nitrogen. CA and DG regions were homogenized in 500 µl ice-cold PBS+Protease Inhibitor Cocktail (PIC) (Sigma Aldrich). Homogenized samples were centrifuged for 5 min, 1500 rpm, 4°C, and supernatant was removed. 1% PFA (fresh prepared from 16% Formaldehyde/Thermo Scientific, Article 28906) was added to the pellet and incubated for 15 min at 22°C in rotation. To stop the reaction, glycine (2.5 M) was added and incubated 5 min in rotation. Samples were centrifuged 6 min, 1500 rpm, 4°C. Samples were washed with PBS+PIC two times. The fixed cells were collected by 5 min, 1500 rpm at 4°C and supernatant was removed. Ice-cold lysis buffer 0,1% SDS+ PIC was added. Each pellet was transferred to a 4°C chilled Dounce homogenizer with tight fitting pestle (Type A), and the samples were homogenized on ice. Cell lysis was confirmed under a microscope. The chromatin of the lysed cells was sheared for 3×10 min in the Bioruptor (30 s pulse, 30 s pause, and high power). The samples were centrifuged for 10 min, 13000 rpm, 4°C. Samples were snap frozen in liquid nitrogen and stored at −80°C until following ChIP-seq experiments.

ChIP-seq libraries were generated from ChIP-DNA using a custom Illumina library type on an automated system (Apollo 342, Wafergen Biosystems/Takara, Active Motif). ChIP-seq libraries were sequenced on Illumina NextSeq 500 as single-end 75bp reads (Active Motif).

#### In vitro primary hippocampal neuron samples

Wild-type or virally transduced primary hippocampal neurons were fixed and collected for subsequent ChIP-seq experiments according to the instructions of the servicing company (Active Motif). Briefly, cells were fixed by adding 1/10 volume formaldehyde solution (11% Formaldehyde, 0.1 M NaCl, 1 mM EDTA pH=8.0, 50 mM HEPES pH=7.9) directly to the medium and incubated at room temperature (RT) with agitation. Fixation was stopped by adding 1/20 volume 2.5 M Glycine solution (Roth, Germany) and incubating for 5 min at RT. Cells were scraped from the plates and transferred into conical tubes, and centrifuged at 800 rpm at 4°C for 10 min. Cells were resuspended and washed twice in chilled PBS-Igepal solution (0.5% Igepal CA-360 (Sigma) in 1X PBS pH 7.4 (Gibco)), and a third time in PBS-Igepal-PMSF (phenylmethylsulfonyl fluoride) solution (0.5% Igepal, 1 mM PMSF (Sigma) in 1X PBS pH 7.4). The supernatant was completely removed and cell pellets were snap frozen in liquid nitrogen and stored at −80°C until following ChIP-seq experiments.

ChIP-seq libraries were generated from ChIP-DNA using a custom Illumina library type on an automated system (Apollo 342, Wafergen Biosystems/Takara). ChIP-seq libraries were sequenced on Illumina NextSeq 500 as single-end 75bp reads (Active Motif).

### RELACS ChIP-seq

Restriction enzyme-based labeling of chromatin *in situ* (RELACS) was used for H3K27ac and H3K4me3 ChIP-sequencing (25). DIV11 primary hippocampal neurons with FOXG1 or luciferase KD were fixed in 1% formaldehyde for 15 min. The reaction was stopped with 125 mM glycine for 5 min, followed by two-time DPBS +PIC washes. Cell nuclei were isolated following the NEXSON protocol (treatment time 20 s) and permeabilized with 0.5% SDS (26). Chromatin was digested *in situ* using five units of restriction enzyme CviKI-1 (NEB, R0710S) every 100.000 nuclei and RELACS custom barcodes (4 bp UMI + 8 bp RELACS barcode) were ligated to end-repaired and A-tailed chromatin using components from NEBNext Ultra II DNA Library Prep Kit for Illumina (NEB) (25).

The barcoded chromatin fragments were extracted by sonication for 5 min using these parameters: peak power 105 W, 2% duty factor, 200 cycles/burst (Covaris microtubes (520185), Covaris E220 Focused-ultrasonicator). A single immunoprecipitation (IP) reaction for all pooled samples was carried out on SX-8G Compact IP-Star platform (Diagenode) following Arrigoni et al. (25). Immunoprecipitated chromatin was used for NGS library preparation (NEBNext Ultra II DNA Library Prep Kit for Illumina, NEB). Libraries were sequenced at the Max Planck Institute of Immunology and Epigenetics using HiSeq 3000 (Illumina) as 75 bp reads.

### ATAC-seq

ATAC-seq was performed on DIV11 primary hippocampal neurons after FOXG1 or luciferase KD following the protocol of Buenrostro et al., 2015 (27). Briefly, cells were washed with PBS and detached by 5 min 0.05% trypsin incubation. Dissociation was stopped with 10% FBS and cells were collected and centrifuged for 5 min at 500 rpm at 4°C. After one time wash with ice-cold PBS, cells were counted and separated into 5×10^5^cells/replicate/condition. Cells were resuspended in lysis buffer (10 mM Tris-HCl pH 7.4, 10 mM NaCl, 3 mM MgCl2, 0.1% Igepal Ca-630) and immediately centrifuged for 10 min at 500 rpm at 4°C. Transposition, PCR amplification and DNA library preparation were done using Nextera DNA library prep kit according to the manufacturer’s instructions (Illumina). DNA was purified and eluted using MinElute PCR purification kit (Qiagen). Libraries were sequenced at the Max Planck Institute of Immunology and Epigenetics using HiSeq 3000 (Illumina) (paired-end 75bp reads).

### Co-Immunoprecipitation (Co-IP) and Immunoblotting

Tissue or N2A cells were lysed in lysis buffer (300 mM NaCl, 20 mM Tris, 1 mM EDTA, 0.5% nonidet-40 (NP-40), pH 7.4) supplemented with protease inhibitor (Sigma-Aldrich) and incubated for 30 min on ice, triturating every 10 min 20 times. After a 10 min centrifugation at 13000 rpm, the supernatant was collected. The salt and NP-40 concentrations were brought down to 100 mM and 0.15%, respectively, using equilibration buffer (20 mM Tris, 1mM EDTA, pH 7.4). Protein concentrations were determined with Bradford reagent (Bio-Rad). 10% input was saved and equal amounts of protein were used for all Co-IPs. Protein G Dynabeads (10004D, ThermoScientific) were washed once with Co-IP buffer (100 mM NaCl, 20 mM Tris, 1 mM EDTA, 0.15% NP-40 pH 7.4), and incubated with Co-IP antibodies (rabbit HDAC2, Cell signaling; rabbit NEUROD1, Abcam) or control IgG antibodies (rabbit IgG kch-504-250, Diagenode) at 4°C overnight. Antibody-coupled beads were washed once with co-IP buffer before co-IP. Cell lysates were blocked with Protein G Dynabeads for 1 h at 4°C for preclearing, subsequently transferred to antibody-coupled beads, and incubated overnight at 4°C in rotation. Antibody-coupled beads were washed 3 times with Co-IP buffer before they were resuspended in 1X Laemmli sample buffer. Protein-antibody were eluted by incubating the beads at 70°C for 10 min at 550 rpm. 10% input and the complete Co-IP sample were used for immunoblotting.

Co-IP samples were loaded on 8% SDS-polyacrylamide gels and run at 120 V for 1.5 h in Tris-Glycine running buffer. Proteins were transferred to PVDF membranes (Trans-blot Turbo Transfer Pack, Bio-Rad) using the Trans-blot Turbo Transfer System (Bio-Rad) following the manufacturer’s instructions. Membranes were blocked with 5% BSA in TBS (Tris buffered saline) for 1 h, and incubated overnight with primary FOXG1 antibody detecting both FOXG1 monomers and dimers (Rabbit polyclonal FOXG1, Active Motif) diluted in 5% BSA in TBST (Tris buffered saline with 0.1% Tween-20). Membranes were washed 3 times with TBST and incubated with secondary antibody anti-rabbit HRP (1:10000 dilution in 5% BSA in TBST,) for 1 h. Membranes were washed 2 times with TBST and 1 time with TBS before being developed using SuperSignal™ West Femto Maximum Sensitivity Substrate (Thermo Scientific). Membranes were imaged using LAS ImageQuant System (GE Healthcare).

### Tissue sections, in situ hybridization and immunostaining

PFA fixed brains were embedded in TissueTec (SAKURA) cut in 14 μm sections and mounted on SuperFrost Plus Microscope slides (Thermo Scientific). Probes for *in situ* hybridization were made by cloning PCR products into pGemTeasy (Promega). 1 µg of the linearized plasmid was transcribed *in vitro* using NTP labelling mix and T7 or sp6 RNA Polymerase, followed by purification with mini Quick spin RNA columns (Roche). Probes were diluted in hybridization buffer in 1:500 or 1:1000 ratio and incubated on the sections overnight at 68°C. After washing and blocking in lamb serum in MABT buffer (5X MAB, 0.1% Tween 20), sections were incubated with Anti-DIG-AP (Roche) at 4°C overnight. After washing, sections were developed with NBT/BCIP (Roche) overnight and mounted. Bright field images were obtained using an Axioplan M2 microscope (Zeiss). Immunofluorescence was performed as described previously (28, 29) using antibodies against SATB2, CTIP2 and ZBTB20. Images were obtained using a primary magnification of 20x, and stitched together to ensemble the complete hippocampus using an Axioplan M2 fluorescent microscope (Zeiss) equipped with an Apotome.2 module.

### DiI tracing

Brains from 7-week old mice were dissected and immersion-fixed in 4% paraformaldehyde in 0,1 M phosphate buffer for 2-3 days and horizontal tissue sections of 2-3 mm thickness were cut by a razor blade. Small crystals of DiI (1,1’-dioctadecyl-3,3,3’,3’-tetramethylindocarbocyanine perchlorate) mounted on the tip of a glass micropipette were placed into different regions of the hippocampal formation under visual control. The brain sections were then stored for 5-6 weeks in the fixative solution at room temperature and in the dark to allow the labelling of fiber pathways. Following this time, sections were washed in PB and sliced into 100 µm thick vibratome sections and mounted on glass slides for fluorescence microscopy. Digital images were made using either Axioplan M2 (Zeiss) or ApoTome 1 (Zeiss).

### N2A cell culture

Mouse neuroblastoma cell line, Neuro-2a (N2A) were cultured in Dulbecco’s modified Eagle’s medium (DMEM, ThermoScientific) supplemented with 10% fetal bovine serum (FBS, ThermoScientific), 1% non-essential amino acids (NEAA, ThermoScientific), 1% L-Glutamine, and 1% Penicillin, Streptomycin, and Neomycin (PSN, ThermoScientific). Cells were maintained at 37°C, 95% relative humidity and 5% CO_2_. Cells were seeded in 6 well plates and transfected with Lipofectamine LTX (ThermoScientific) adding a total amount of 2.5 μg plasmid according to the instructions of the manufacturer.

### Dual Luciferase Reporter Assay

50.000 N2A cells were seeded to 24-well plates overnight and double-transfected the following morning with 2.5 µg pMirGlo luciferase reporter constructs and pLenti-III overexpression plasmid DNA using Lipofectamine LTX (Thermo Scientific) reagent. pMirGlo reporters contained inserts with exemplary *Ncald* or *Ldb2* regulatory regions with or without targeted ablation of Fkh or bHLH/E-Box motifs. pLenti-III overexpression constructs contained inserts with a sequence coding for the intact FOXG1 protein (FOXG1 OE) or a protein bereft of the Fkh-binding domain (Foxg1 ΔFKH) (30).Medium was changed after 24 h. 48 h after transfection, cells were washed once with 1X PBS and incubated with 1X Passive Lysis Buffer (Promega) at room temperature for 20 min, gently agitating. Lysates were loaded in triplicate to 96-well microtiter plates and luminescence was measured in a GloMax 96 microplate luminometer (Promega). Gene expression was measured by firefly luciferase activity (LAR II; Promega) normalized against the *Renilla* luciferase signal (Stop & Glo; Promega), which served as a control. Background noise was filtered out of both signals prior to calculation by subtracting luminosity measured before enzyme injection. All promoter constructs were tested with at least three biological replicates.

### Luciferase reporter sequences

#### Ldb2

TTAACATGATGTTAATTATTTGTAAATTGTATGTTTGGATACACATTACTTGAGAGGCATATGTTATAT TCAA

#### Ldb2, Fkh deleted

TTAACATGATGTTAATTAGTAAATTGTATGGGATACACATTACTTGAGAGGCATATGTTATATTCAA

#### Ldb2, E-box delete

TTAACATGATGTTAATTATTTGTAAATTGTATGTTTGGATACACATTACTTGAGAGGCAGTTATATTCA A

#### Ncald

GGAACAATGCAACCAAATATGGACATTCTATTTAGAAACAGTAATGTTAGCATAACAAGGGAGCTTAAAAAAAGAAAAAGCTAAATAAATATTAAACAGATGGGTGGTCAAATGTTGTTTTTCTTTTTAAGCACAAAGCCTTAATCTAAAGCCAAGCAG

#### Ncald, Fkh deleted

GGAACAATGCAACCAAATATGGACATTCTAAGCAGTAATGTTAGCATAACAAGGGAGCTTAAAAAAAGAAAAAGCTTATTCAGATGGGTGGTCAAATGTTGTTTTTCTTTTTAAGCACAAAGCCTTAATCTAAAGCCAAGCAG

#### Ncald, E-box delete

GGAACAATGCAACCAAATATGGACATTCTATTTAGAAACAGTAATGTTAGCATAACAAGGGAGCTTAAAAAAAGAAAAAGCTAAATAAATATTAAACAGGGTGGTCAAATGTTGTTTTTCTTTTTAAGCACAAAGCCTTAATCTAAAGCCAAGCAG

#### Ncald, Fkh & E-box delete

GGAACAATGCAACCAAATATGGACATTCTAAGCAGTAATGTTAGCATAACAAGGGAGCTTAAAAAAAGAAAAAGCTTATTCAGGGTGGTCAAATGTTGTTTTTCTTTTTAAGCACAAAGCCTTAATCTAAAGCC AAGCAG

### Statistics

GraphPad Prism software (version 9.1.1) was used for statistical analyses. Two-way ANOVA followed by Tukey multiple comparisons was used for dual luciferase assay analyses. Unpaired t-test was used for testing qRTPCR results. Fisher’s exact test was used to test the enrichment of DEGs in clusters using GeneOverlap package in R Bioconductor(v. 3.8).

### Artwork

The graphical abstracts were illustrated using Biorender.com.

### Bioinformatics, Data repository and analyses of public databases

The “Differential Search” tool of the Allen Brain Atlas (31) was used to define field specific gene expression. The respective subfield was set as target structure and the other fields as contrast structure. The emerging list was visually inspected with ISH datasets to confirm selective expression in the field as well as overall expression. Assignment to a field was performed according to clearly visible ISH signal. From this list we compiled the subset of genes that were tested in qRTPCR analysis.

The sequencing data from RNA-, ATAC-, RELACS-, and wildtype ChIP-seq were processed with snakePipes (v. 1.1.1) (32). Relevant parameters used for each experiment and summary QC are available at https://github.com/ipekakol/FOXG1. Mapping was performed on mouse genome build mm10 (GRCm38). For ChIP-Seq and ATAC-seq, high quality and uniquely mapping reads were retained (mapq > 5). RELACS custom barcodes were designed with integrated UMI, so duplicate removal was performed using UMITools57 (33), while a standard deduplication was applied for ATAC-seq reads. For ChIP-seq and ATAC-seq data, snakePipes also provided candidate peak regions using MACS2 (v. 2.2.6) using default parameters.

Differential expression analysis for RNA-seq was done using DESeq2 (v. 1.22.1) on count matrices output from snakePipes (featureCounts, v. 1.6.4). A linear model controlling for batch effects (e.g., ∼batch + treatment or ∼ batch + condition) was used and apeglm log_2_(Fold Change) shrinkage was applied.

Differential ChIP-seq and ATAC analyses were performed on consensus peak sets, coverage was computed using multiBamSummary (deeptools) (34) and differential regions identified via csaw (v. 3.13).

The sequencing data from FOXG1 and NEUROD1 ChIP-seq after NEUROD1 orFOXG1 KD were uploaded to the Galaxy web platform, and the public server at usegalaxy.eu was used to analyze the data (35). Same parameters were applied for quality control and mapping as the snakepipes analyses. Peaks were called using MACS2 callpeak (Galaxy Version 2.1.1.20160309.6). Coverage was computed using multiBamSummary, and bam files were normalized by bamcompare and bigwigcompare (deeptools, Galaxy Version 3.3.2.0.0) (34). Differential binding regions of NEUROD1 were computed using DiffBind (Galaxy Version 2.10.0) (36).

All metaprofiles and heatmaps of ChIP- and ATAC-seq signals were generated with deeptools (Galaxy Version 3.3.2.0.0). All ChIP- and ATAC-seq peaks were annotated and visualized using ChIPSeeker (Galaxy Version 1.18.0).

GimmeMotifs (v. 0.13.1 and v. 0.16.0) was used for motif enrichment and differential motif analysis. The transcription factor affinity prediction (TRAP) method was used to calculate the affinity of FOXG1 and NEUROD1 for the peak sequences at +/− 200 bp flanking intronic and intergenic FOXG1-peak summits retrieved from *in vivo* and *in vitro* WT FOXG1 ChIP-seq (37). GO enrichment and differential GO-term analyses were performed using clusterProfiler (R, v. 3.10.1) (38).

Visualizations were produced in Galaxy, R (v. 4.1) and Python (v. 3.6). Heatmaps were plotted using heatmap2 (Galaxy Version 3.0.1). Volcano plots were plotted using EnhancedVolcano (BioConductor, v. 3.13). Violin plots were plotted using ggplot2 (v. 3.3.5). Venn diagrams were plotted using ggVenn and VennDiagram packages. Fisher’s exact test was applied using the GeneOverlap package (Bioconductor, v. 3.8) (39).

### Hipposeq

Hipposeq (23) was accessed at http://hipposeq.janelia.org/ choosing the dorsal-ventral survey of hippocampal principal cells. Dorsal and ventral populations of CA1, CA2 and CA3 were chosen and the gene names were entered as a list using the standard settings of the database, and the count matrix was normalized as transcripts per million (TPM). The set of genes with significant increase or decrease in expression from our RNA-seq were intersected with the TPM normalized count matrix from hipposeq. The genes found in the intersection were clustered using hierarchical clustering on the row-wise Z score.

### STRING Database

STRING Database (24) was used to explore known and predicted protein-protein interactions of FOXG1 in *Mus musculus*. Data from experiments and databases were used with medium confidence (0.4) for the interaction scores. Nodes annotated interactor proteins, while edges connected interacting proteins. Line thickness indicated the confidence of interaction. The interactome layout was reorganized in Cytoscape (v 3.8.2) to depict direct interactions of FOXG1 with HDACs ad HATs.

## Results

### Reduced FOXG1 expression alters gene transcription in the adult mouse hippocampus *in vivo* and primary hippocampal neurons *in vitro*

Since (i) part of the phenotypic impairments of FOXG1-syndrome affect hippocampal functions, (ii) FOXG1 haploinsufficiency has not been studied at the chromatin level in the context of the hippocampus, and (iii) we only have limited insights into FOXG1 function in later developmental stages of the hippocampus, we decided to use the postnatal hippocampus with reduced expression of FOXG1 as a model to unravel molecular mechanisms used by FOXG1 to control transcription at the chromatin level.

*Foxg1* was found to be expressed in the DG as well as granule neurons residing in the CA fields *in vivo.* The *Foxg1*-heterozygous (Foxg1^cre/+^) hippocampus contained all fields but was generally smaller compared to wild type (WT) (**Fig. S1A**). The expression of mature granule cell marker genes (CTIP2, SATB2, ZBTB20) did not reveal obvious differences between the genotypes (**Fig. S1B**) and the tri-synaptic neuronal network of the hippocampus was similarly unchanged (**Fig. S1C**). Despite a grossly normal appearing Foxg1^cre/+^ hippocampus, we revealed transcriptomic changes compared to WT *in vivo,* as well as *in vitro* after shRNA-mediated FOXG1 knockdown (KD) compared to control. RNA-sequencing (RNA-seq) of Foxg1^cre/+^ and WT hippocampus revealed 174 differentially expressed genes (DEGs) (105 increased, 69 decreased) within a threshold of log_2_ Fold Change (LFC) ±log_2_(1.5) and an adjusted p-value of ≤0.05 (**Fig. 1A**). Genes with increased expression enriched mainly for gene ontology (GO)-terms angiogenesis or inflammation, conferring limited insight into FOXG1 functions specifically in the hippocampus. In contrast, genes with decreased expression enriched in GO-terms related to nervous system development or function, including synaptogenesis, in the top 10 significant hits (**Fig. 1B**). We also generated RNA-seq data after shRNA-mediated KD of FOXG1 in cultivated primary hippocampal neurons *in vitro*, because this experimental set up would allow biochemical manipulation to explore FOXG1 functions in a comparably less complex cell context. The RNA-seq after FOXG1 KD and luciferase KD as respective control retrieved 2626 DEGs (821 increased, 1805 decreased) using the same thresholds (**Fig. 1C**). GO-terms enriched in increased DEG reflected forebrain development, whereas neuronal differentiation and synaptogenesis dominated DEG with decreased expression (**Fig. 1D**). We used publicly available data on region-specific transcriptomes of the hippocampus (23) and revealed that reduced levels of FOXG1 *in vivo* not only affected the DG (14), as previously described, but also the CA fields. Transcripts with increased expression upon FOXG1 reduction clustered with the CA3/4-field-enriched marker genes, whereas decreased expression was recognized for CA1-field-specific transcripts (**Fig. S2A, B**).

**Figure 1:**
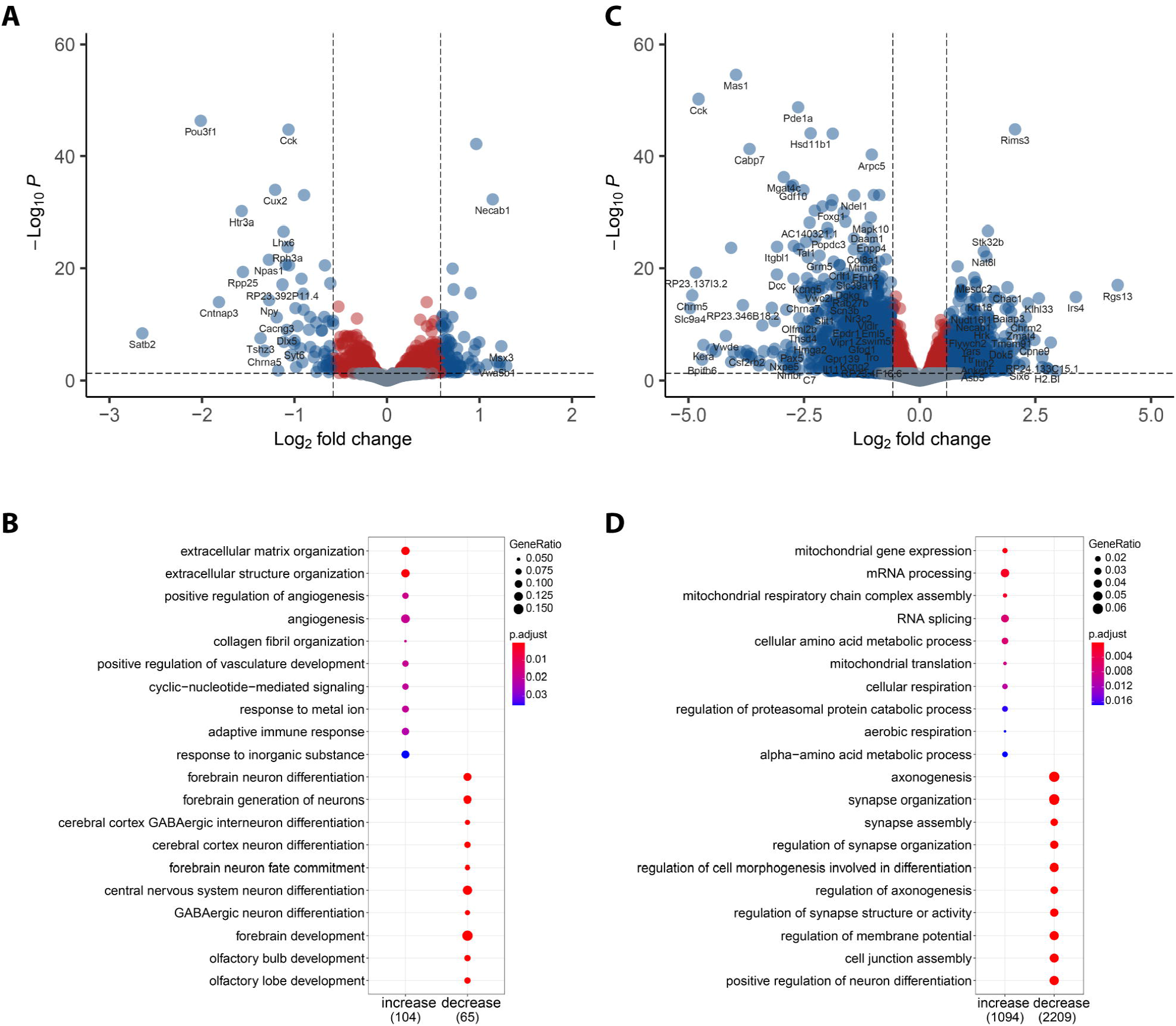
*Foxg1* haploinsufficiency alters the transcriptome of the mouse hippocampus *in vivo* and *in vitro.* **(A)** Volcano plot displaying differentially expressed genes (DEGs) between hippocampus samples from *Foxg1*^cre/+^ and *Foxg1*^+/+^ (n=2). The y-axis corresponds to the adjusted p-value and the x-axis displays the log_2_Fold Change (log_2_FC). Grey: Transcripts with insignificant adjusted p-values (p>0.05). Red: Transcripts with differential expression of less than ±log_2_(1.5). Blue: Transcripts with differential expression of more than ±log_2_(1.5). Positive log_2_FC represents increase, negative log_2_FC decrease in expression upon *Foxg1*-haploinsufficiency. **(B)** Dotplot of top 10 functionally enriched GO-terms in the fraction of increased or decreased DEGs in the adult *Foxg1^cre/+^* hippocampus. Total number of increased or decreased genes are on the x-axis, dot size displays the gene ratios and color indicates significance level of the enrichment. Threshold for enrichment analysis is adjusted to p < 0.01. **(C)** Volcano plot displaying DEGs upon reduction of FOXG1 in primary neurons (n=5). The y-axis corresponds to the adjusted p-value, and the x-axis displays the log_2_FC value. Color code as in A. **(D)** Top 10 functionally enriched GO-terms in DIV7 primary neurons upon KD of FOXG1. Representation as in B.

Together, these analyses indicated that reduced expression of FOXG1 decreased the hippocampus size without further major impact on the gross structure, but with a defined altered transcriptional program *in vivo* and *in vitro*, affecting, amongst others, neuronal differentiation and synaptogenesis.

### FOXG1 binds chromatin in intronic and intergenic regions *in vivo* and *in vitro*

To correlate the observed transcriptional changes upon FOXG1 reduction with its presence and thus action at the chromatin level, we explored the binding of FOXG1 to the DNA. We performed ChIP-seq, again both *in vivo* (from 6-week old hippocampus tissue) and *in vitro* (from primary hippocampal neurons). The comparison of both data sets was intended to explore whether insights from *in vitro* cultivated cells could be used, at least partially, to understand FOXG1 functions *in vivo*.

As our transcriptome data had shown that FOXG1 reduction affected both DG and CA fields, we used hippocampus tissue subdivided into DG or CA fields for *in vivo* ChIP-seq. FOXG1 was bound to unique loci in both the DG and CA data sets, but we also detected shared chromatin regions. The percentage and the pattern of the distribution of FOXG1 peaks within the genome was similar in both DG and CA fields, and we observed a high degree of overlap of functional GO-terms of genes associated with FOXG1 peaks (**Fig. S2C-F**). As we did not concentrate on region-specific FOXG1 functions in further experiments, but rather aimed to study general chromatin functions of FOXG1, we merged DG and CA FOXG1 peaks for *in vivo* ChIP-seq data sets for subsequent analyses.

We addressed next (i) the nature of the binding regions of FOXG1 in hippocampus tissue and in cultivated hippocampal neurons, and (ii) the degree of overlap in the binding pattern in the two different cellular sources. For both *in vivo* and *in vitro* data sets, to the largest extent, FOXG1 peaks were distributed genome-wide to distal intergenic and intronic regions, and a smaller fraction to promoters (**Fig. 2A**). We used the transcriptional start sites (TSS) as reference points to analyze localization of FOXG1 peaks at promoters. GO-term analysis of regions where FOXG1 peaks allocated to specific genes revealed a large overlap between *in vivo* and *in vitro* data sets, with neuronal differentiation and synaptic functions represented within the top 10 enriched terms (**Fig. 2B**). This was generally in line with the GO-term signature based on the transcriptional changes observed upon reduced expression of FOXG1 (**Fig. 1B, D**). Using k-means (with k=5) revealed similarities between *in vivo* and *in vitro* (**Fig. 2C**). Two clusters had FOXG1 peaks at or near the TSS, two other clusters indicated a less discrete localization of FOXG1, and one cluster was not enriched in FOXG1 binding. Cluster-wise GO-term analysis of the FOXG1 cistrome at promoter regions showed in the first two clusters that genes with FOXG1-associated peaks affected axono- and synaptogenesis both *in vivo* and *in vitro* (**Fig. 2D, F**). Genes affecting mitochondrion organization and non-coding RNA biology were represented in cluster 4 *in vivo* and cluster 3 *in vitro*. Thus, largely, cluster-wise GO-term analysis retrieved overlapping classifications *in vivo* and *in vitro*.

**Figure 2:**
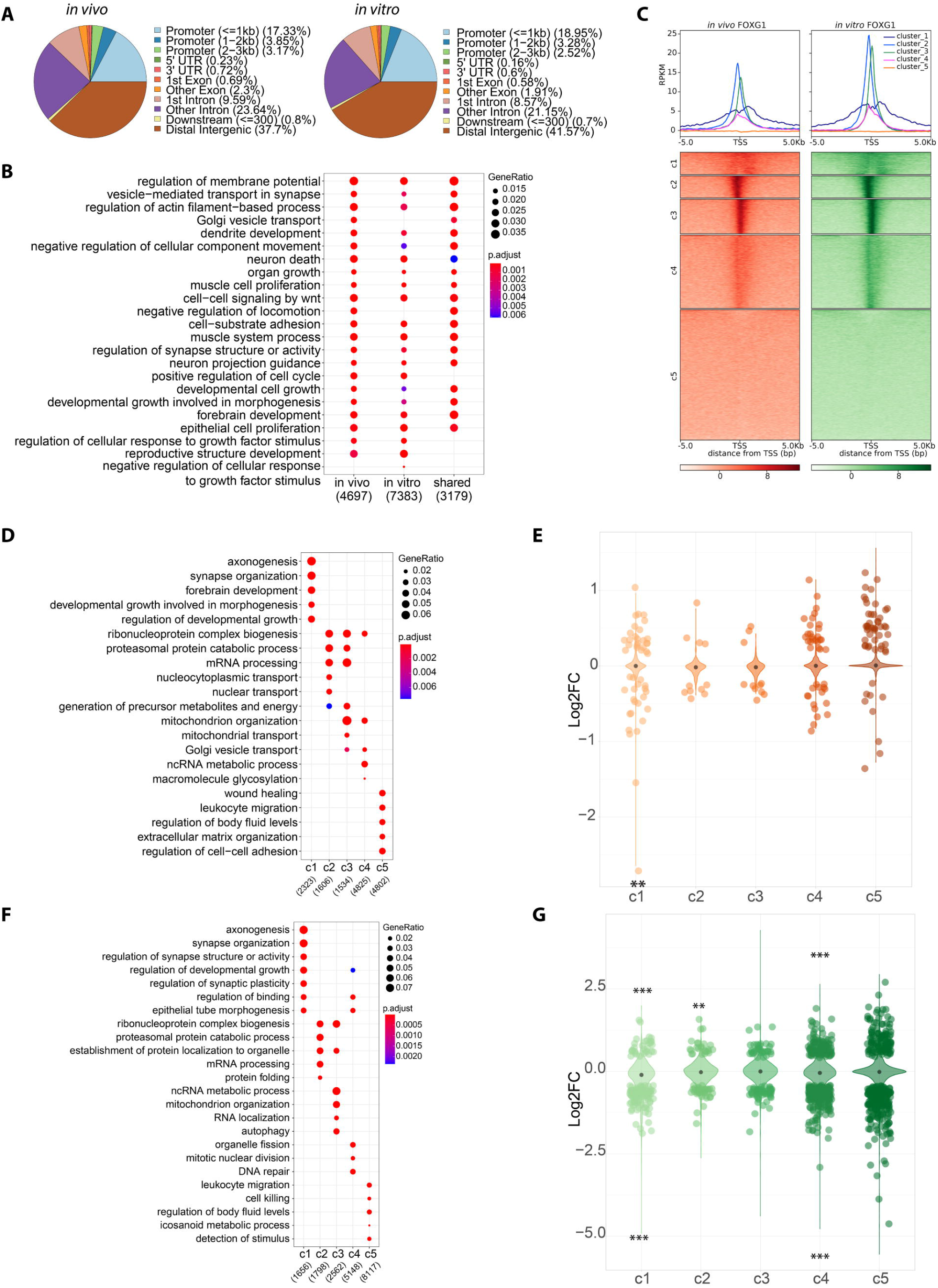
FOXG1 enriches at intergenic, intronic and promoter regions in the hippocampus *in vivo* and *in vitro*. **(A)** Pie charts depicting the genomic distribution of FOXG1 peaks *in vivo* (left), and *in vitro* (right) and showing FOXG1 enrichment in distal intergenic (brown), intronic (pink, purple), and promoter (blue) regions. **(B)** K-means clustering (k=5) of FOXG1 enrichment *in vivo* (orange) and *in vitro* (green) found 5 Kb up-/down-stream of transcription starts sites (TSS) of protein coding genes. Data is normalized by sequencing depth and input control. The metaprofiles (top) show the average reads per kilo base per million mapped reads (RPKM) of each cluster. **(C)** Dotplot of functional GO-enrichment analysis showing the profiles for *in vivo, in vitro*, and their shared FOXG1 peaks. **(D)** Dotplot of cluster-wise GO-enrichment analysis of FOXG1 occupancy *in vivo* (according to clusters shown in A). Gene ratios and adjusted p-values are indicated at the top-right corner and total number of genes per group are on the x-axis. Threshold for enrichment analysis is adjusted to p < 0.01 **(C, D)**. **(E)** Violin plot correlating FOXG1 enrichment in k-means clusters *in vivo* (clusters as in A) and DEGs upon FOXG1 haploinsufficiency. Y-axis corresponds to log_2_FC of gene expression, and x-axis shows clusters. The black dot marks the median of log_2_FC of DEGs in each cluster. **(F)** Cluster-wise GO-enrichment analysis of FOXG1 enrichment *in vitro* (according to clusters shown in A). Dotplot as in D. (**G**) Violin plot correlating FOXG1 enrichment in k-means clusters *in vitro* (clusters as in A) and DEGs upon KD of FOXG1. Representation as in E. **(E, G)** Fisher’s exact test *: p<0.05, **: p<0.01, ***: p<0.001.

We plotted the distribution of DEGs upon reduced expression of FOXG1 in the five different clusters of FOXG1-peak locations for *in vivo* and *in vitro* data sets. We observed DEGs in all 5 clusters, both with increasing and decreasing levels, and statistically significant enrichment of DEGs with either increased or decreased expression in specific clusters (**Fig. 2E, G**).

We concluded that (i) the binding patterns of FOXG1 observed *in vivo* and *in vitro* overlapped largely, (ii) genes in the vicinity of FOXG1 peaks *in vivo* and *in vitro* confer similar functions, and (iii) genes in the vicinity of FOXG1 peaks can increase or decrease in expression upon FOXG1 reduction both *in vivo* and *in vitro*. This high degree in overlap between *in vivo* and *in vitro* cellular resources prompted us to study *in vitro* primary hippocampal neurons in further experimentation, aiming to unravel molecular mechanisms used by FOXG1 to control gene expression at the chromatin level.

### Reduced levels of FOXG1 alters the epigenetic landscape

#### Reduced levels of FOXG1 increase and decrease H3K27 acetylation

FOXG1 ChIP peaks localized at putative enhancer and promoter regions. We next aimed to confirm this localization and to elucidate concomitantly whether FOXG1 impacts transcription through epigenetic mechanisms. We addressed these questions by using quantitative ChIP-seq of H3K27ac. Wild type and FOXG1 KD *in vitro* primary hippocampal neurons were used and correlative mapping of FOXG1- and H3K27ac-peaks confirmed co-occurrence of FOXG1 with H3K27ac, thus presence of FOXG1 at enhancers in hippocampal neurons (**Fig. 3A**). As we hypothesized, KD of FOXG1 in primary hippocampal neurons indeed altered H3K27ac levels, both increasing and decreasing H3K27ac. In three out of four k-means clusters, FOXG1 and H3K27ac peaks co-occurred. H3K27ac levels did not change upon KD in cluster 1, increased in cluster 2, decreased in cluster 3, and cluster 4 did not have any noticeable levels. The H3K27ac signal in the H3K27ac-stable cluster 1 mapped genome-wide to around 60% at promoter-assigned sites, whereas the regions with increasing (cluster 2) and decreasing (cluster 3) H3K27ac levels localized respectively to only 30% and 10% to promoters, but mapped predominantly to intergenic and intronic regions (**Fig. 3B**). GO-terms of annotated genes correlating with FOXG1/H3K27ac-peaks in the four clusters enriched for synapse organization and axonogenesis, as well as signal transduction (**Fig. 3C**). Differential transcription factor (TF)-binding-motif analysis within 200bp flanking the peak summits in the four clusters showed that cluster 1 regions enriched for cell-cycle control TFs, including HES proteins or the E2F family, in addition to CXXC1. Cluster 2 specifically enriched for Forkhead (Fkh) TFs, and clusters 2, 3 and 4 enriched for basic helix-loop-helix (bHLH) and GATA motifs (**Fig. 3D**). DEGs upon FOXG1 KD distributed to all four clusters, where cluster 1 (unchanged high levels of H3K27ac) was significantly enriched in genes with increased and decreased expression, and cluster 3 (decreasing levels of H3K27ac) contained significantly more genes with decreased expression, in accordance with reduced levels of H3K27ac (**Fig. 3E**).

**Figure 3:**
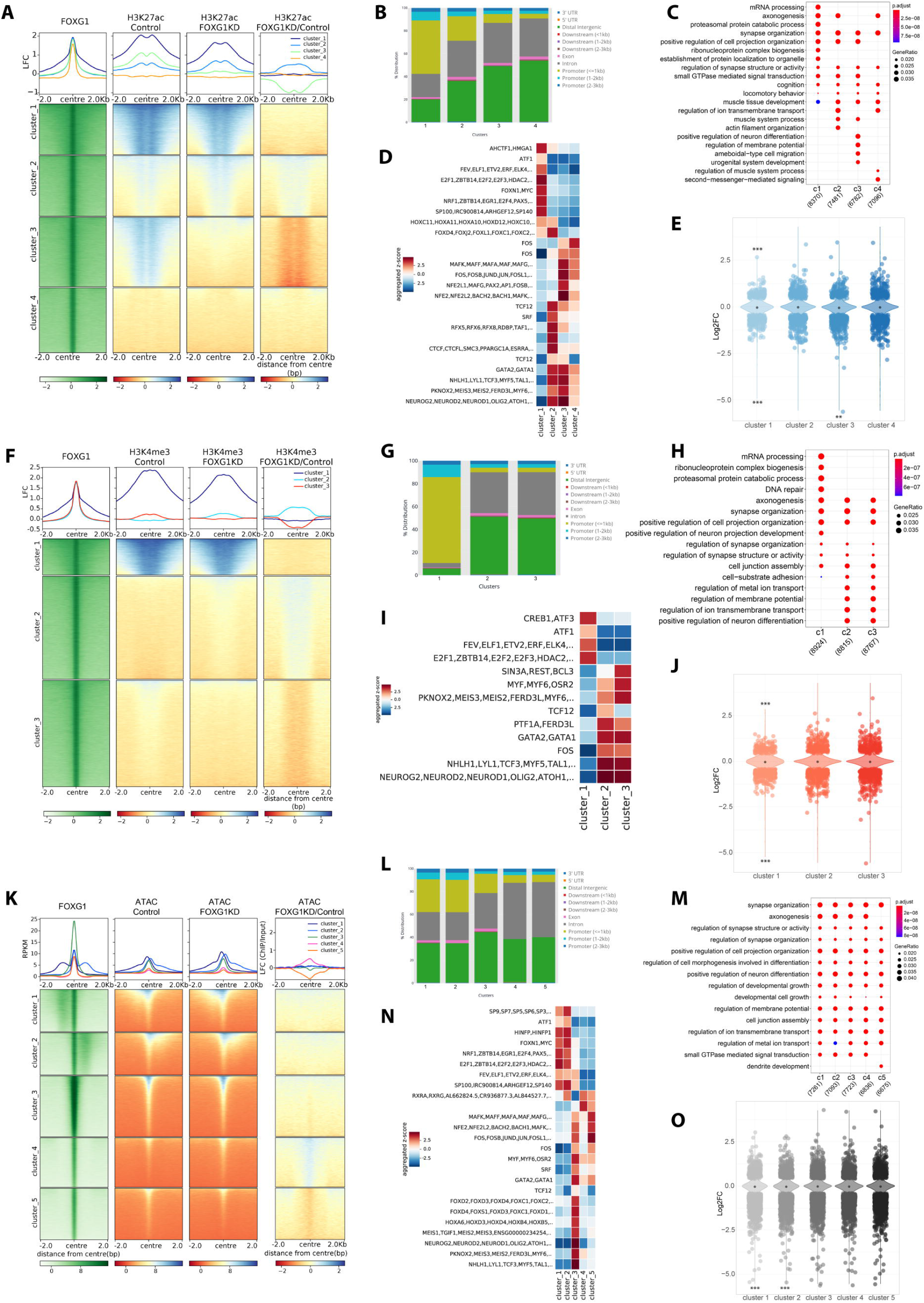
Reduced levels of FOXG1 alter the epigenetic landscape. **(A)** Heatmap of k-means clustered (k=4) H3K27ac enrichment at 2Kb up-/down-stream of FOXG1 peak summits (left, green) in control and FOXG1 KD conditions. Data is normalized by sequencing depth and input control as log_2_ (ChIP/Input) for FOXG1, H3K27ac control and H3K27ac FOXG1 KD data. The difference between FOXG1 KD and control conditions is calculated from RPKM normalized bigwig files as log_2_ (FOXG1 KD/Control). The metaprofiles (top) show the mean log_2_FC (LFC) of each cluster. **(B)** Genomic distribution of H3K27ac enrichment at FOXG1 peaks, according to k-means clusters from A, displayed as a stacked bar graph. **(C)** Enriched GO-terms for the respective k-means clusters as shown in A. Scales of gene ratios and adjusted p-value are at top-right corner, and total number of genes per cluster are on the x-axis. Threshold for enrichment analysis was adjusted to p < 0.01. **(D)** Transcription factor (TF)-binding differential motif analysis according to the clusters of H3K27ac enrichment at FOXG1 binding regions as shown in A. **(E)** Violin plot depicting the distribution of DEGs upon FOXG1 KD at k-means clusters of H3K27ac enrichment at FOXG1 peak as shown in A. Y-axis corresponds to log_2_FC of gene expression, and x-axis shows the four clusters. The black dot marks the median of log_2_FC of DEGs in each cluster. **(F)** Heatmap of k-means clustered (k=3) H3K4me3 enrichment 2Kb up-/down-stream of FOXG1 peak summits (left, green) in control and FOXG1 KD conditions. Data representation as in A. **(G)** Genomic distribution of clustered H3K4me3 enrichment at FOXG1 peaks represented as a stacked bar graph as shown in F. **(H)** GO-term analysis comparing enrichment of H3K4me3 enrichment at FOXG1 peaks according to k-means clusters shown in F. Representation as in C. **(I)** TF-binding differential motif analysis of the three k-means clusters of H3K4me3 enrichment at FOXG1 binding regions. **(J)** Violin plot depicting the distribution of DEGs upon FOXG1 KD at the three k-means clusters (according to F) of H3K4me3 enrichment at FOXG1 peaks. Representation as in E. **(K)** Heatmap of k-means clustered (k=5) ATAC enrichment 2Kb up-/down-stream of FOXG1 peak summits (left, green) in control and FOXG1 KD conditions. Data representation as in A. **(L)** Genomic distribution of ATAC enrichment according to the five k-means clusters shown in K at FOXG1 peaks displayed as a stacked bar graph. **(M)** GO-term comparison between the five k-means clusters (K) of ATAC enrichment at FOXG1 peaks. Representation as in C. **(N)** TF-binding differential motif analysis of the five clusters of ATAC enrichment at FOXG1 binding regions. **(O)** Violin plot depicting DEGs upon FOXG1 KD at five k-means clusters of ATAC enrichment at FOXG1 peaks (according to K). Representation and statistics as in E. Fisher’s exact test, *: p<0.05, **: p<0.01, ***: p<0.001.

We also analyzed only those regions that significantly gained or lost H3K27ac upon FOXG1 KD. This confirmed that (i) regions with altered H3K27ac levels localized predominantly to introns and intergenic regions (**Fig. S3A, B**), (ii) DEGs significantly enriched for genes with increased expression and increased H3K27ac, and vice versa for decreased expression and decreased H3K27ac (**Fig. S3C**), (iii) axonogenesis and neuron projection extension enriched as GO-terms with genes showing altered expression and H3K27ac levels (**Fig. S3D**), and (iv) specific TF motifs enriched in regions gaining or loosing H3K27ac upon FOXG1 reduction (**Fig. S3E**).

We concluded that FOXG1 controls transcription by altering H3K27ac levels, predominantly at enhancers, which might impact binding of other TFs, including members of the bHLH family.

#### Reduced levels of FOXG1 alter H3K4 trimethylation

We used H3K4me3 ChIP-seq that is found at both promoters and enhancers, and investigated whether FOXG1 controls transcription by impacting this epigenetic modification. We used k-means clustering with k=3 to explore the H3K4me3 profile at FOXG1 binding sites in FOXG1 KD cells compared to controls. Two clusters displayed mild changes in H3K4me3 levels. Cluster 2 had slightly increased levels of H3K4me3, and cluster 3 had slightly decreased levels (**Fig. 3F**). The majority of the H3K4me3 peaks were found at promoters for the cluster 1 as expected, whereas the regions gaining or losing H3K4me3 upon FOXG1 KD mapped with 90% to intronic or intergenic regions (**Fig. 3G**). Among enriched GO-terms were synaptogenesis and axonogenesis (**Fig. 3H**). TF motifs enriched in the respective clusters again contained CXXC1 in the stable cluster, as well as bHLH and GATA TF in the clusters with altered H3K4me3 (**Fig. 3I**). DEGs upon FOXG1 reduction were distributed to all clusters, and significantly more genes either with increased or decreased expression were found in cluster 1, which had the strongest H3K4me3 levels (**Fig. 3J**). Focused analyses of the regions with significant gain or loss of H3K4me3 upon FOXG1 KD showed that (i) these regions were classified mainly as promoters (**Fig. S4A, B**), (ii) the distribution of DEGs was significantly enriched for genes with decreased expression and decreased H3K4me3 (**Fig. S4C**), and axon extension enriched as GO-term within the genes that lost H3K4me3 alongside other terms indicative of neuronal differentiation (**Fig. S4D**).

We concluded that FOXG1 alters transcription by altering H3K4me3 levels, but the extend seemed milder compared to H3K27ac. Altered H3K4me3 levels predominantly localized at enhancers, also impacting binding sites for bHLH family members. Therefore, FOXG1 uses epigenetic mechanisms at enhancers, and impacts function of other TFs.

#### Reduced levels of FOXG1 increase and decrease chromatin accessibility

Presence of FOXG1 at regulative regions, i.e. enhancers, its action towards altering epigenetic marks, and the prediction that other TF might be affected at these locations, prompted us to complete our epigenetic survey by analyzing opening and closing of chromatin regions upon FOXG1 KD. ATAC-seq revealed that FOXG1 peaks mapped indeed to opened and closed chromatin regions, and k-means clustering associated the cluster 4 with gain of accessibility and clusters 3 and 5 with loss of accessibility upon FOXG1 KD (**Fig. 3K**). As with the other epigenetic parameters, the altered clusters mainly contained regions between genes or in introns (**Fig. 3L**). Similarly, GO-terms enriched for axono- and synaptogenesis, as well as signal transduction (**Fig. 3M**), and the TF motifs at the FOXG1 peaks included CXXC1, E2F, HES, bHLH and GATA TF (**Fig. 3N**). DEGs distributed again to all clusters, with clusters 1 and 2 showing significant differences in the distribution between decreased or increased genes (**Fig. 3O**).

The focused analyses of the regions that significantly gained or lost accessibility upon FOXG1 KD affirmed impact at enhancer regions (**Fig. S5A, B**). Interestingly, compared to H3K27ac and H3K4me3, the FOXG1 peaks distributed more sharply around the center of the peak in the regions that changed in accessibility (**Fig. S3A, S4A, S5A**). All clusters contained DEGs (**Fig. S5C**) and synaptogenesis enriched as a GO-term within the genes that gained access, while signal transduction was enriched within genes that lost access (**Fig. S5D**). Importantly, bHLH and GATA TF motifs enriched also in regions gaining access upon FOXG1 reduction (**Fig. S5E**).

Summarizing the entire epigenetic survey, we conclude that FOXG1-bound chromatin localized to enhancer, and to a smaller extend to promoter regions. The epigenetic alterations studied upon FOXG1 KD affected H3K27ac and H3K4me3 levels as well as chromatin accessibility, whereby both gain and loss of marks/accessibility was observed. Affected genes classified as regulating axono- and synaptogenesis. Epigenetic changes impacted regions that can be bound by TFs, among which bHLH family members were associated in all three altered chromatin contexts. Thus, FOXG1 is a TF with multi-modal activities at the chromatin level, one of which is alteration of the epigenetic landscape.

### HDAC inhibition reverts a fraction of FOXG1 transcriptional changes

To elucidate mechanistically how FOXG1 alters the epigenetic landscape, we decided to focus on its impact on H3K27ac because of the predominant localization of FOXG1 at enhancers. We observed both increasing and decreasing levels of H3K27ac upon FOXG1 KD, but reasoned that because FOXG1 does not have HAT or HDAC activity itself, it might regulate access of these enzymes to the chromatin. Indeed, the publicly available network of proteins interacting with FOXG1 (STRING database) suggested association with both HATs (EP300) and HDACs (SIRT1) (**Fig. 4A**). Our own interactome study after overexpression of FOXG1 in N2A cells ((30)) indicated as well potential association with HDACs (HDAC1, 2, SIRT1) (**Fig. 4B**). Confirming these indications, FOXG1 co-immunoprecipitated with HDAC2 from hippocampal tissue (**Fig. 4C**). We thus hypothesized that using epigenetic drugs impacting histone acetylation could be a potential way to develop novel therapeutic strategies for the human disease.

**Figure 4:**
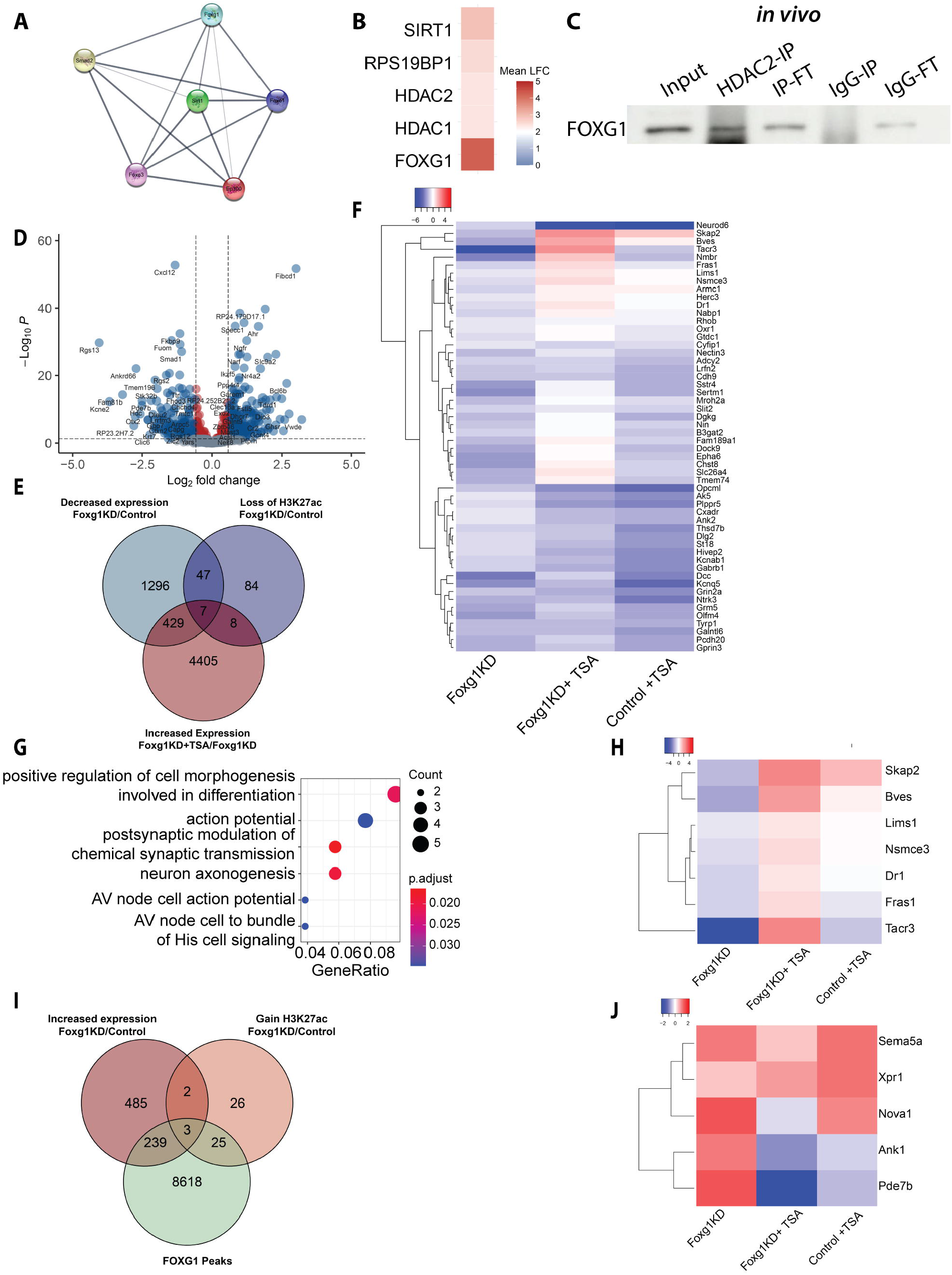
FOXG1 influences HDAC function. **(A)** STRING DB predictions of FOXG1-interacting proteins shows FOXG1 interaction with HATs and HDACs. Each node represents an interactor protein. Known interactions between proteins are retrieved from curated databases and experimental data. Line thickness indicates the strength of data support. **(B)** Heatmap of enriched HATs and HDACs upon FOXG1 pull-down after its overexpression in N2A cells according to (30). **(C)** Immunoblots after co-IP demonstrating interaction between HDAC2 and FOXG1 dimer in hippocampal tissue from adult animals. n=1 **(D)** Volcano plot showing 956 DEGs of DIV11 hippocampal neurons upon TSA treatment alongside FOXG1 KD. Differential expression is analyzed for TSA/DMSO conditions upon FOXG1 KD. DEGs obtained are intersected with DEGs upon FOXG1 KD. DEGs upon TSA treatment under shCtrl conditions are removed to exclude FOXG1-independent effects of HDAC inhibition. The y-axis corresponds to the adjusted p-value, and the x-axis displays the log_2_FC. Color code and thresholds as represented as in Fig. 1A. **(E)** DEGs assuming FOXG1-HDAC interaction according to repression model. Venn diagram shows the intersection of DEGs with decreased expression and decreased H3K27ac upon reduced levels of FOXG1, and DEGs with increased expression upon TSA treatment after FOXG1 KD. **(F)** Heatmap of 54 genes at the intersection of reduced H3K27ac and gene expression upon FOXG1 KD resulting from D. Scale shows log_2_FC upon respective conditions shown in the x-axis. **(G)** GO-term enrichment analysis shows the biological processes affected in the 54 genes according to repression model. **(H)** Heatmap of 7 genes at the intersection of reduced H3K27ac and gene expression upon FOXG1 KD, and rescued upon TSA treatment. Scale as in F. **(I)** DEGs assuming FOXG1-HDAC interaction according to recruitment model. Venn diagram demonstrates the intersection of FOXG1 peaks, DEGs with increased expression and gain of H3K27ac upon reduced levels of FOXG1. **(J)** Heatmap of 5 genes at the intersection of gain of H3K27ac and increased gene expression upon Foxg1KD assuming the recruitment model resulting from I. Scale as in F.

To test this hypothesis, we treated primary hippocampal neurons with the broad HDAC inhibitor Trichostatin A (TSA) in the condition of FOXG1 KD (shFoxg1) or control (shControl) and assessed the corresponding transcriptional alterations using RNA-seq. Bioinformatics analyses comprised of (i) the identification of DEGs from shFoxg1/shControl, (ii) intersection of this DEG set with DEGs from TSA/DMSO in shFoxg1, and (iii) exclusion of DEGs in TSA/DMSO in shControl to eliminate FOXG1-independent effects of HDAC inhibition. This approach retrieved 956 DEGs that changed expression upon TSA treatment in FOXG1 KD (**Fig. 4D**).

We next tested for two hypotheses as to how FOXG1 could affect HDACs to influence gene transcription. Firstly, FOXG1 could prevent binding of HDACs to the chromatin (repression model, **Fig. S6A**), or could otherwise recruit HDACs to the chromatin (recruitment model, **Fig. S6B**). The repression model is independent of FOXG1-binding to the chromatin and predicts that reduced levels of FOXG1 lead to reduced levels of H3K27ac (shFoxg1/Ctrl decreased H3K27ac: 156 loci) and concomitant transcriptional decrease (shFoxg1/Ctrl decreased expression: 1779 DEG). Upon HDAC inhibition with TSA, we expect transcriptional increase of genes (shFoxg1+TSA/shFoxg1 increased expression: 4849 DEG) regulated through the repression model. Intersection of the respective gene fractions showed that 54 genes decreased in expression alongside decreased H3K27ac levels (**Fig. 4E, F**). GO-term analysis returned functional terms for synaptic functions and axonogenesis (**Fig. 4G**). 7 genes of the 54 fulfilled all criteria for the repression model, i.e. increasing upon TSA inhibition alongside FOXG1 KD (**Fig. 4E, H**). Among these target genes regulated in a setting following the repression model were *Lims1* and *Fras1*, which are linked to epilepsy and neurite growth, processes that are affected by FOXG1 mutation.

The recruitment model predicts that reduced levels of FOXG1 correlate with increased H3K27ac levels (shFoxg1/Ctrl increased H3K27ac: 56 loci) and concomitantly with increased transcription (shFoxg1/Ctrl increased expression: 729 DEG) at FOXG1-binding regions (FOXG1 peaks: 9066 loci) (**Fig. 4I**). In this case, TSA treatment would not alter transcription upon reduced FOXG1 levels, as inhibition of the HDACs does not occur bound to the chromatin. Five genes had increased expression and H3K27ac levels upon FOXG1 reduction, three of which were also bound by FOXG1 (**Fig. 4I, J**). All criteria for the recruitment model was fulfilled by one gene, *Sema5a*, which is an autism-susceptibility gene with a role in axon guidance and synaptogenesis.

We concluded from these findings that HDAC inhibitor treatment might be worth further exploration as potential drug to alleviate transcriptional alterations upon reduced FOXG1 expression. In regard to the mechanism, FOXG1 seems to affect both functional repression and recruitment of HDACs, as shown for a small subset of target genes.

### FOXG1 and NEUROD1 act in concert to regulate axono- and synaptogenesis genes in hippocampal neurons

A further perception of our study of the epigenetic alterations upon reduced FOXG1 expression was that other TF, especially bHLH family proteins acted in synergistic or antagonistic fashion to FOXG1. For some Fkh- and bHLH-domain proteins pioneering functions and thus the potential to alter the epigenetic landscape have been assigned (40–42). We therefore explored the hypothesis that FOXG1 acted together with bHLH TF. To this aim, we determined, using bioinformatics predictions, the affinities of TF binding to FOXG1 peaks at intronic and intergenic regions that we retrieved from FOXG1 ChIP-seq *in vivo* and *in vitro* as predominant localization of FOXG1. Indeed, at these putative enhancer regions containing FOXG1 peaks, we observed not only enrichment of Fkh TF motifs, but also surprisingly a significant co-occurance of bHLH TF motifs (**Fig. S7A, B**). Importantly in the context of neuronal functions, we identified the bHLH TF NEUROD1 binding motif as significantly enriched at FOXG1 peaks (**Fig. 5A**), both *in vivo* and *in vitro*. We also noted that motif analysis generally ranked bHLH motifs at a higher significance level than Fkh motifs (**Fig. 5B**). Moreover, Fkh/bHLH motifs mainly co-occurred in intergenic regions and introns, while the Fkh motif was also enriched in proximal promoter regions (**Fig. 5C**).

**Figure 5:**
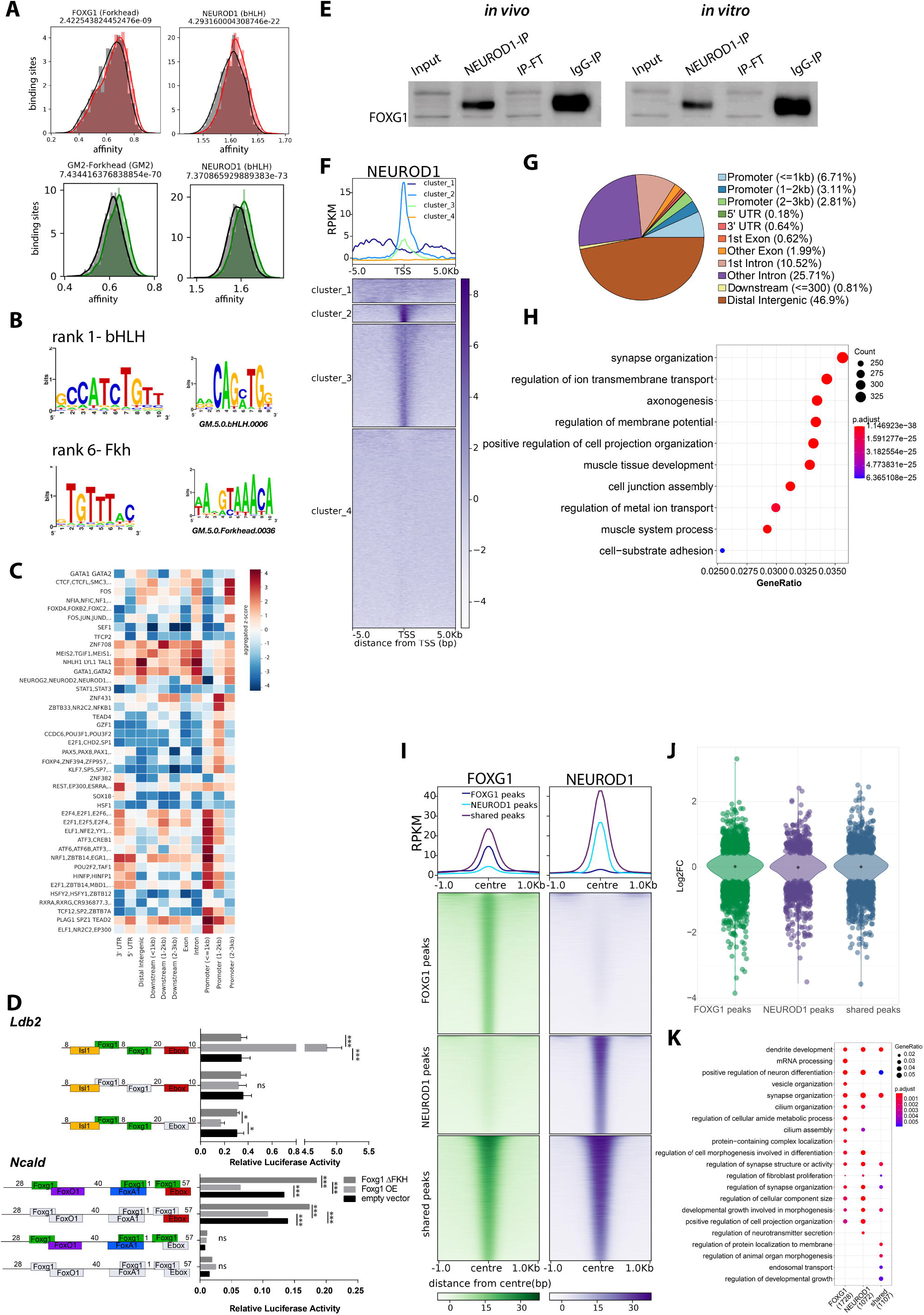
FOXG1 and NEUROD1 interact and share binding sites at genes implicated in neuronal functions. **(A)** Affinities of FOXG1 and NEUROD1 to bind sequences in a range from +/− 200 bp flanking FOXG1 peak summits at intronic and intergenic regions *in vivo* (red), and *in vitro* (green) as predicted by TRAP motif affinity analysis. FOXG1 and NEUROD1 have both significantly high affinities at FOXG1 binding sites compared to dinucleotide-shuffled background control. Y-axis shows the binding site numbers, while x-axis indicates the calculated affinity scores. **(B, C)** Enriched TF binding motifs at +/− 200bp of FOXG1 summits include bHLH and Fkh motifs *in vivo* (B) and *in vitro* (C). **(D)** Luciferase reporter assays showing FOXG1 activity on regulatory regions of two target genes containing Fkh- and bHLH/E-box motifs in direct vicinity, *Ncald* (top) and *Ldb2* (bottom). Regulatory regions containing WT or modified sequences are illustrated next to the graphs along the y-axis. Deleted motifs are shown in grey, while present binding motifs are shown for FOXG1 (green), FOXO1 (purple), FOXA1 (blue), bHLH/E-box motif (red), and ISL1 (yellow). Relative luciferase activity upon overexpression of full length FOXG1 (Foxg1 OE, light grey), FOXG1 with deleted Fkh domain (Foxg1 ΔFKH, dark grey), or empty vector (control, black) are shown on the x-axis. Two-way ANOVA with Tukey’s multiple comparisons: *: p<0.05, **: p<0.01, ***: p<0.001, ns: non-significant. **(E)** Representative immunoblot after NEUROD1 co-IP using anti-FOXG1 antibody shows that monomeric FOXG1 and NEUROD1 interact *in vivo* in adult hippocampus (left), and *in vitro* in primary hippocampal neurons (right). n=1 **(F)** K-means clustering (k=4) of NEUROD1 enrichment *in vitro* at 5 Kb up- /down-stream of TSS of protein coding genes. Data is normalized by sequencing depth and input control. The profiles (top) show the average RPKM of each cluster. **(G)** Pie chart of genomic distribution of NEUROD1 peaks *in vitro* shows NEUROD1 enrichment mainly in distal intergenic (brown) and intronic (pink, purple) regions, and to a lesser extent, promoter (blue) regions. **(H)** GO-term enrichment analysis of NEUROD1 peaks. Scales of gene ratio and adjusted p-value reported at the top right. Total number of genes per cluster are on the x-axis. Threshold for enrichment analysis was adjusted to p < 0.01. **(I)** Heatmap showing FOXG1 (green) and NEUROD1 (purple) binding sites clustered into shared and unique (FOXG1_unique, NEUROD1_unique) regions. Data normalization and metaprofiles (top) as in F. **(J)** Violin plot demonstrates the distribution of DEGs upon reduced levels of FOXG1 at unique and shared clusters of FOXG1 and NEUROD1 binding sites according to I. y-axis corresponds to log_2_FC of gene expression, and x-axis shows clusters. The black dot marks the median of log_2_FC of DEGs in each cluster. **(K)** Dotplot shows the top 10 terms of biological processes enriched in unique and shared clusters of DEGs corresponding to FOXG1 and NEUROD1 peaks.in. p-value threshold as in H.

We next used luciferase reporter assays to investigate FOXG1/bHLH co-activity exemplarily at regulative regions of two DEGs, *Ldb2* and *Ncald,* that both contain Fkh and bHLH (called E-box) motifs in direct vicinity (**Fig. 5D**). *Ldb2* increased and *Ncald* decreased upon FOXG1 overexpression, and this opposing response gave the possibility to explore diverse mechanisms of FOXG1-mediated transcriptional control. FOXG1 activation of *Ldb2* transcription was abolished upon deletion of its Fkh protein domain, and was dependent on the FOXG1-binding motifs in the reporter construct. Presence of the bHLH/E-box binding sequence seemingly prevented a repressive FOXG1 function. FOXG1 repression of *Ncald* transcription was also dependent on the presence of its Fkh domain, and the deletion of the Fkh domain increased reporter transcription. Deletion of the binding sites for FOXG1 in the regulatory region still resulted in the same response of the reporter transcription as observed for the unmodified construct. The deletion of the bHLH/E-box binding site silenced the regulatory region, independent of the presence or absence of Fkh motif.

We concluded that for these two regulatory regions, FOXG1 modulated the bHLH-mediated transcriptional regulation. Repression of transcription depended on the Fkh domain but was independent of FOXG1 binding sites on the DNA. In contrast, activation of transcription depended on the Fkh domain and on FOXG1 binding sites, whereas both FOXG1 repressive or activating functions depended on the presence of the E-box motif. Thus, the transcriptional responses of FOXG1 in concert with bHLH appeared to be context-dependent and multimodal. The observation of a crosstalk of FOXG1 with bHLH TFs prompted us to explore this in more detail with a focus on FOXG1 and NEUROD1. We decided on NEUROD1, as (i) its binding motif significantly enriched at FOXG1 peaks, (ii) it is a pro-neuronal bHLH transcription factor expressed in mature hippocampal neurons, with essential roles in neuronal development and function (43, 44), similar to FOXG1, and (iii) it has been associated with pioneering activity (41). Further, co-immunoprecipitation showed that FOXG1 and NEUROD1 precipitated together, and that they might therefore act in concerted manner at the chromatin level (**Fig. 5E**).

We next aimed to confirm FOXG1/NEUROD1 co-occurance on the chromatin level by determining the cistrome of NEUROD1 in primary hippocampal neurons. The NEUROD1 ChIP-seq profile, centered around the TSS, showed enrichment of NEUROD1 peaks at these sites but also up- and downstream (**Fig. 5F**). Genome-wide distribution profiles indicated that the majority of peaks localized at intergenic and intronic regions (**Fig. 5G**). GO-terms enriched for genes with NEUROD1 peaks contained axono- and synaptogenesis, similar to what we observed for FOXG1 peak distribution (**Fig. 5H**). We therefore compared FOXG1 and NEUROD1 ChIP-seq profiles, which returned clustered genes bound by both (shared fraction) and also genes that only enriched binding of one of the TFs (FOXG1, NEUROD1 fraction) (**Fig. 5I**), with enrichment of genes implicated in synapse organization and axonogenesis (**Fig. S7C**). Interestingly, for all three clusters, including NEUROD1 unique peaks, we observed DEGs upon FOXG1 reduction with both increased and decreased expression (**Fig. 5J**). This finding consolidated our viewpoint that FOXG1 and NEUROD1 act in concert, in chromatin-dependent and -independent fashions, the latter of which is displayed by altered expression of NEUROD1 uniquely-bound genes upon FOXG1 KD. Again, axono- and synaptogenesis were enriched GO-terms for the gene fraction with both FOXG1- and NEUROD1-bound peaks and altered expression levels upon FOXG1 KD (**Fig. 5K**). We plotted the observed epigenetic changes upon reduced expression of FOXG1 to the regions that were shared or enriched for either FOXG1 or NEUROD1 (**Fig. S8A**). Only a small fraction of genes with peaks for FOXG1, NEUROD1, or both displayed epigenetic alterations upon FOXG1 reduction. The moderate alterations were confined to altered chromatin accessibility, whereas genome-wide H3K27ac or H3K4me3 levels were affected very little.

We concluded that FOXG1 and NEUROD1 co-occurred at the chromatin level and that they have a remarkable overlapping set of target genes conferring neuronal functions. Alteration of the epigenetic landscape at regions bound by both TF was moderate, indicating another mechanism by which these two key instructors affect transcriptional programs important for axono- and synaptogenesis.

### FOXG1 and NEUROD1 act in a concerted manner, both at the chromatin level and prior to chromatin binding

We next aimed to define whether FOXG1 and NEUROD1 would act in a concerted manner, or up-/downstream from each other. This question is of interest to place either factor in the driver seat regarding neuronal differentiation. Thus, to analyze the nature of the FOXG1/NEUROD1 crosstalk in more detail, we knocked down either FOXG1 or NEUROD1 in primary hippocampal neurons and assessed the binding profile of the other respective TF using ChIP-seq. We clustered the peak regions according to the shared fraction and regions that enriched predominantly for either FOXG1 or NEUROD1 (**Fig. 6A**). Within the shared regions, FOXG1 binding was reduced upon NEUROD1 KD, and vice versa, NEUROD1 reduced upon FOXG1 KD, excluding a clear hierarchy in this crosstalk.

**Figure 6:**
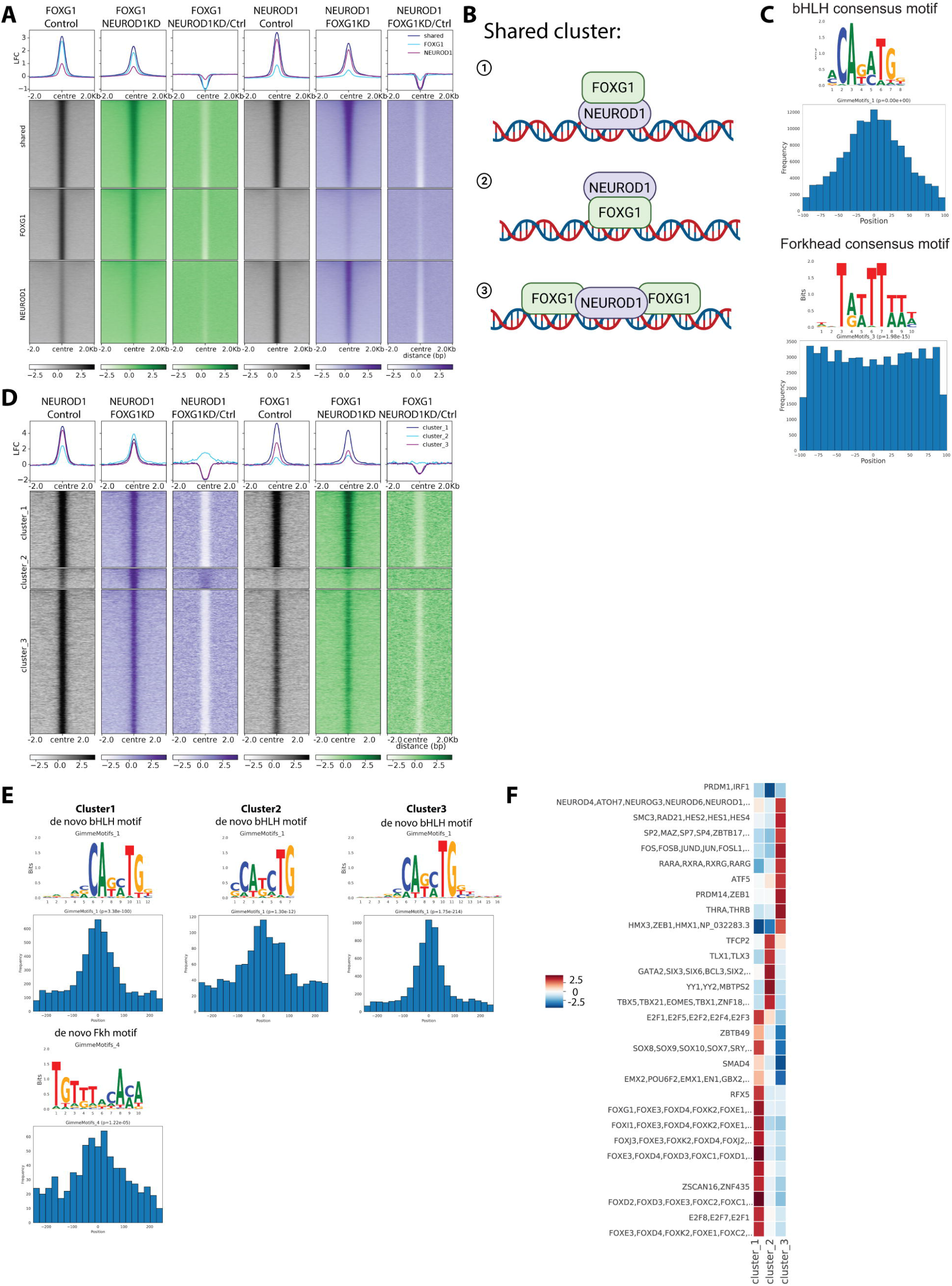
FOXG1 and NEUROD1 act in concert rather than up- or downstream from each other. **(A)** Heatmap of FOXG1 (green) and NEUROD1 (purple) enrichment clustered into unique and shared regions under control (grey), NEUROD1 KD, and FOXG1 KD conditions. Data is normalized by sequencing depth and input control as log_2_ (ChIP/Input). The difference of FOXG1 KD/Control and NEUROD1 KD/Control are calculated from RPKM normalized bigwig files as log_2_ (NEUROD1 KD/Control or FOXG1 KD/Control). The metaprofiles (top) show the average log_2_FC (LFC) of each cluster. **(B)** Cartoon of binding modes that classify for categorization as shared binding sites. 1: NEUROD1 binds to its bHLH/E-box motif at the chromatin and indirectly brings in FOXG1. 2: FOXG1 binds to its Fkh motif at the chromatin and indirectly brings in NEUROD1. 3: Binding sites of NEUROD1 and FOXG1 co-occur near a respective peak center (example depicts NEUROD1 as peak center). **(C)** Positional preference plots and motif logos of bHLH (top) and Fkh (bottom) motifs at FOXG1/NEUROD1 shared regions retrieved from de novo motif analysis. **(D)** Heatmap of k-means clustered (k=3) NEUROD1 (purple) and FOXG1 (green) enrichment 2Kb up-/down-stream of differential NEUROD1 binding sites retrieved from DiffBind analysis between FOXG1 KD and control conditions. Data representation as in A. **(E)** Positional preference plots and motif logos of bHLH (top) and Fkh (bottom) motifs at sites with significant alteration of NEUROD1 binding upon FOXG1 KD, according to the three clusters from D, retrieved from de novo motif analysis. **(F)** Heatmap showing differential TF-binding motif analysis clustered at differential NEUROD1 binding sites as shown in B.

Shared binding regions could reflect binding through one of the respective factors and indirect presence at the site of the other, or usage of adjacent binding sites (**Fig. 6B**). To resolve these binding paradigms, we determined the distribution of the respective other binding motif in the vicinity of either the FOXG1 or NEUROD1 peaks within the shared regions. The majority of NEUROD1 motifs distributed directly in the center of the FOXG1 peaks (**Fig. 6C**). This finding suggests that NEUROD1 binds most likely directly to chromatin and brings in FOXG1 alongside, thus FOXG1 is indirectly recruited to chromatin at the majority of these sites. In contrast, FOXG1 motifs at NEUROD1 peaks were distributed broadly around and flanking the center of NEUROD1 peaks (**Fig. 6C**). Thus, at NEUROD1 peaks FOXG1 could recruit NEUROD1 to one extend, but also flanking, cooperative binding would be possible. We concluded that rather than having a clear hierarchy within the crosstalk of both TF, they are acting multi-modal and cooperative.

Surprisingly, FOXG1 KD decreased NEUROD1 presence at NEUROD1 sites (hardly enriched for FOXG1). And vice versa, NEUROD1 KD decreased FOXG1 presence at FOXG1 sites (hardly enriched for NEUROD1). We concluded that either factor might be needed before or during recruitment but not for stable binding as well.

We next analyzed the changes of NEUROD1 and FOXG1 binding upon the respective KDs, with a focus on those regions that showed differential binding of NEUROD1 upon FOXG1 KD. Regions presenting differential binding of NEUROD1 clustered into three main profiles: Cluster 1 had strong enrichment for NEUROD1 and FOXG1, cluster 2 had moderate enrichment for NEUROD1 and lower enrichment of FOXG1, and cluster 3 had strong enrichment for NEUROD1 and moderate enrichment of FOXG1 (**Fig. 6D**). Upon FOXG1 KD, binding of NEUROD1 within clusters 1 and 3 (high and moderate levels of FOXG1) reduced. Interestingly, cluster 2, with little enrichment of FOXG1, had increased binding of NEUROD1 after FOXG1 KD. NEUROD1 KD also altered FOXG1 presence at loci within clusters 1 and 3, but not in cluster 2.

We concluded that binding of NEUROD1 and FOXG1 at clusters 1 and 3 is cooperative, either at one binding site (cluster 3) or at adjacent sites (cluster 1), as supported by the distribution of the binding motifs in regard to the peak center (**Fig. 6E**).Cluster 2 targets in contrast are bound mainly by NEUROD1, and at these sites FOXG1 presence interferes with NEUROD1 binding. In support of this interpretation is the cluster-wise analysis of enriched binding motifs. Cluster 2 enriched for example for GATA and TBX TF, but neither for Fkh or bHLH/E-box motifs (**Fig. 6F**). We speculate that NEUROD1 can thus be recruited to chromatin by other than Fkh TF, but that FOXG1 is a competing NEUROD1 binding partner impacting the cluster 2 genomic regions.

## Discussion

The data presented in this study enlighten for the first time in a comprehensive manner diverse and multi-modal functions of FOXG1 at the chromatin level in mature neurons. Despite FOXG1 being recognized as a key transcription factor for telencephalic development and neuronal function, insights into the mechanism underlying transcriptional regulation are sparse. We here show that (i) FOXG1 acts both as repressor and activator, (ii) localizes predominantly to enhancer regions, (iii) alters the epigenetic landscape, (iv) affects directly HDAC functions, and (v) acts in concert with NEUROD1 to instruct transcriptional programs necessary for axono- and synaptogenesis.

FOXG1 is generally considered to be a TF with repressive function (6, 45), but recent data support pleiotropic and context-dependent functions. For example, non-nuclear functions of FOXG1 encompass posttranscriptional regulation (21) and functions in mitochondria (22). Further, different signaling pathways control nuclear or cytoplasmic localization of FOXG1 (46). Surprisingly, chromatin-related functions of FOXG1 are only partly understood, but also seem to be diverse in their nature. For example, FOXG1 exerts transcriptional regulation in ternary protein complexes, hampering FOXO/SMAD transcriptional activators to bind to *Myc*-target regions (47). ZBTB18, a zinc-finger protein binding to E-box core motifs, was proposed as a repressor that was cooperatively recruited with FOXG1 to genes affecting neuronal migration and axonal projections (6). However, the complexity of the FOXG1-syndrome suggests further actions. To fill the gap in knowledge about FOXG1 functions at the chromatin, we report on various high-throughput data sets that extend current views of the multiple mechanisms used by FOXG1 to control gene expression, both as a repressor and an activator. In the hippocampus model system, one important finding is that FOXG1 impacts regulatory genomic regions, mostly enhancers but also promoters, by direct binding and by altering the epigenetic landscape. Thereby, reduced FOXG1 levels correlated with both increased and decreased H3K27ac, H3K4me3, and/or chromatin accessibility, highlighting the diverse context-dependent molecular functions of FOXG1. In regard to FOXG1 impacting H3K27ac, we identified two means of action. On one hand, FOXG1 represses the recruitment of HDACs to chromatin (chromatin-independent function); on the other, it can repress HDAC function at the chromatin (chromatin-dependent function). FOXG1 target genes regulated via HDACs were, for example, *Lims1*, *Fras1*, and *Sema5a,* which are associated with epilepsy, behavioral abnormalities, autism, intellectual disability and impaired synaptogenesis (48–51). These features are also observed in FOXG1-syndrome patients.

In accordance with other studies reporting on FOXG1 cistrome in the cerebral cortex (6, 8), FOXG1 peaks localized at genes that influence maturation and function of neurons in the hippocampus. Axono- and synaptogenesis are functional terms that were enriched in several of our data sets, indicating that FOXG1 influences these processes both by direct presence, by associated epigenetic alterations, and in concert with NEUROD1. Other studies have provided experimental evidence that FOXG1 is important for axonogenesis, synaptogenesis and other features of neuronal differentiation in the hippocampus (18), the retina (52) and the cerebral cortex (6), but lacking the mechanistic layer that our study provides.

Our in-depth analysis of the genome-wide binding pattern of FOXG1 further advances understanding of this TF with highly context-dependent functions. Our data indicate that FOXG1 might have different affinities to chromatin, because FOXG1 reduction seems to affect transcription largely in regions that are moderately bound by FOXG1. Changes in FOXG1, either by loss of function or by increased/decreased abundance, influence proper functioning of the CNS in human patients or animal models (3, 15). Thus, maintaining a critical amount of FOXG1 present at the chromatin seems important for proper neuronal function. Our data suggest that differences in the presence of FOXG1 at respective loci, i.e. loci with less FOXG1 bound, might be particularly vulnerable to association with altered transcription in conditions of reduced FOXG1 presence. Of note, it has been recently shown that other Fkh TF bind DNA as mono- and dimers (53). Despite the fact that our data so far have not the necessary resolution to further analyze action as mono- or dimers, this is an attractive model to explain one aspect of the multi-modal effects of FOXG1 at the chromatin level.

Of particular note is our observation that the FOXG1 presence at the chromatin has a strong correlation with co-occurrence of NEUROD1, a bHLH TF necessary for neuronal differentiation (54, 55). Interestingly, both FOXG1 and NEUROD1 influenced the presence of the other TF at a variety of binding sites in hippocampal neurons.

NEUROD1 has been reported to act as a pioneer factor when its expression was induced in mouse embryonic stem cells (41). In this paradigm of neuronal differentiation, H3K27ac levels increased concomitantly with NEUROD1 activation, and H3K27me3 levels decreased at selected loci. The authors proposed NEUROD1 as a pioneering factor, relieving heterochromatic repression. Our data allow for the assessment as to whether such pioneering function would also be observable in hippocampal neurons, and whether FOXG1 would follow NEUROD1’s pioneering function to access chromatin. We observed a concomitant reduction, but also an increase of binding events for NEUROD1 upon reduced expression of FOXG1. Only a minor fraction of regions that bound altered amounts of NEUROD1 displayed accompanying altered H3K27ac levels upon FOXG1 KD; in 2554 regions that lost NEUROD1 binding, only 23 had significantly lower H3K27ac levels, and in 321 regions that gained NEUROD1, only 1 region had significantly higher H3K27ac levels (data not shown). This observation weakens support for the argument that NEUROD1 acts as a pioneer that alters H3K27ac levels in mature hippocampal neurons upon FOXG1 KD. Because of the reported pioneering potential of NEUROD1, one can also hypothesize that NEUROD1 might act upstream of FOXG1 to regulate its access to the chromatin. However, our data do not support a clear hierarchy of the two factors in hippocampal neurons, but favor the interpretation that they act in concert and both regulate access to each other. Further, our data support the view that, in addition to concerted action through presence at the chromatin, FOXG1 interferes with NEUROD1 binding to chromatin, similar to what has been observed for chromatin accessibility of the SMAD/FOXO TF complexes (47). Thus, FOXG1 and NEUROD1 act together in a chromatin-dependent and - independent manner.

## Conclusion

Together, our data highlight that the multiple modalities of FOXG1 functions found in different cellular compartments, or occurring at posttranscriptional and transcriptional levels, extend to the chromatin level. Here, FOXG1 acts through different epigenetic mechanisms, as well as in concert with other TFs, one of which is NEUROD1. Given that we identified a larger set of TFs, including the Fkh and bHLH families, enriched at FOXG1 bound peaks, the data presented here likely unearth just a small part of the wider epigenetic picture and further studies will add to the complex pattern of different FOXG1 actions. In regard to therapeutic options, direct interference with the litany of TFs acting in concert with or antagonized by FOXG1 would prove challenging. However, considering our observed epigenetic changes upon FOXG1 KD in light of increasing inclusion of epigenetic drugs in clinical trials, our data might highlight an attractive avenue for treatments of FOXG1-syndrome by epigenetic drugs in the future.

## List of abbreviations

ATAC: Assay for Transposase-Accessible Chromatin
BSA: Bovine serum albumin
CA: Cornu ammonis
CNS: Central nervous system
DB: Database
DEG: Differentially expressed gene
DG: Dentate gyrus
DIV: Days in vitro
DMEM: Dulbecco’s Modified Eagle’s Mediu
DMSO: Dimethyl sulfoxide
DPBS: Dulbecco’s phosphate buffered saline
EDTA: Ethylenediaminetetraacetic acid
FBS: Fetal bovine serum
FGF: Fibroblast growth factor
FKH: Forkhead
FOXA1: Forkhead box A 1
FOXG1: Forkhead box G 1
FOXO: Forkhead box O
GABA: Gamma-Aminobutyric acid
GAPDH: Glyceraldehyde-3-phosphate dehydrogenase
GFP: Green fluorescent protein
H3K27: Histone 3 lysine 27
H3K4: Histone 3 lysine 4
H3K79: Histone 3 lysine 79
HAT: Histone acetyl transferase
HBSS: Hank’s buffered salt solution
HDAC: Histone deacetylase
HEPES: N-2-hydroxyethylpiperazine-n’-2-ethanesulfonic acid
IP: Immunoprecipitation
ISH: In stu hybridization
ISL1: Isl lim homeobox 1
KD: Knockdown
KI: Knockin
LFC: Log fold change
MECP2: Methyl cpg binding protein 2
N2A: Neuro2a neuroblastoma
NEAA: Non-essential amino acids
NEUROD1: Neurogenic differentiation 1
NTP: Nucleoside triphosphate
OE: Overexpression
PBS: Phosphate buffered saline
PFA: Paraformaldehyde
PIC: Proteinase inhibitor cocktail
PMSF: Phenylmethylsulfonyl fluoride
PSN: Penicillin-streptomycin-neomycin
QC: Quality control
RELACS: Restriction enzyme-based labeling of chromatin in situ
ROR: Retinoic acid-related Orphan Receptors
RPKM: Reads per kilo base per million mapped reads
RT: Reverse transcription
SATB2: Special AT-rich sequence-binding protein 2
SDS: Sodium dodecyl sulfate
SEM: Standard error of mean
SEMA5A: Semaphorin 5A
SIRT1: Sirtuin 1
SNP: Single nucleotide polymorphism
SOX2: Sry-box transcription factor 2
TBS: Tris buffered saline
TBST: Tris buffered saline with tween
TF: Trasncription factor
TGF: Transforming growth factor
TPM: Transcript per million
TRAP: Transcription factor Affinity Prediction
TSA: Trichostatin A
TSS: Transcription start site
UMI: Unique molecular identifiers
WT: Wild type
ZBTB18: Zinc finger and btb domain containing 18
ZBTB20: Zinc finger and btb domain containing 20

## Legends to supplementary figures

**Figure S1:**
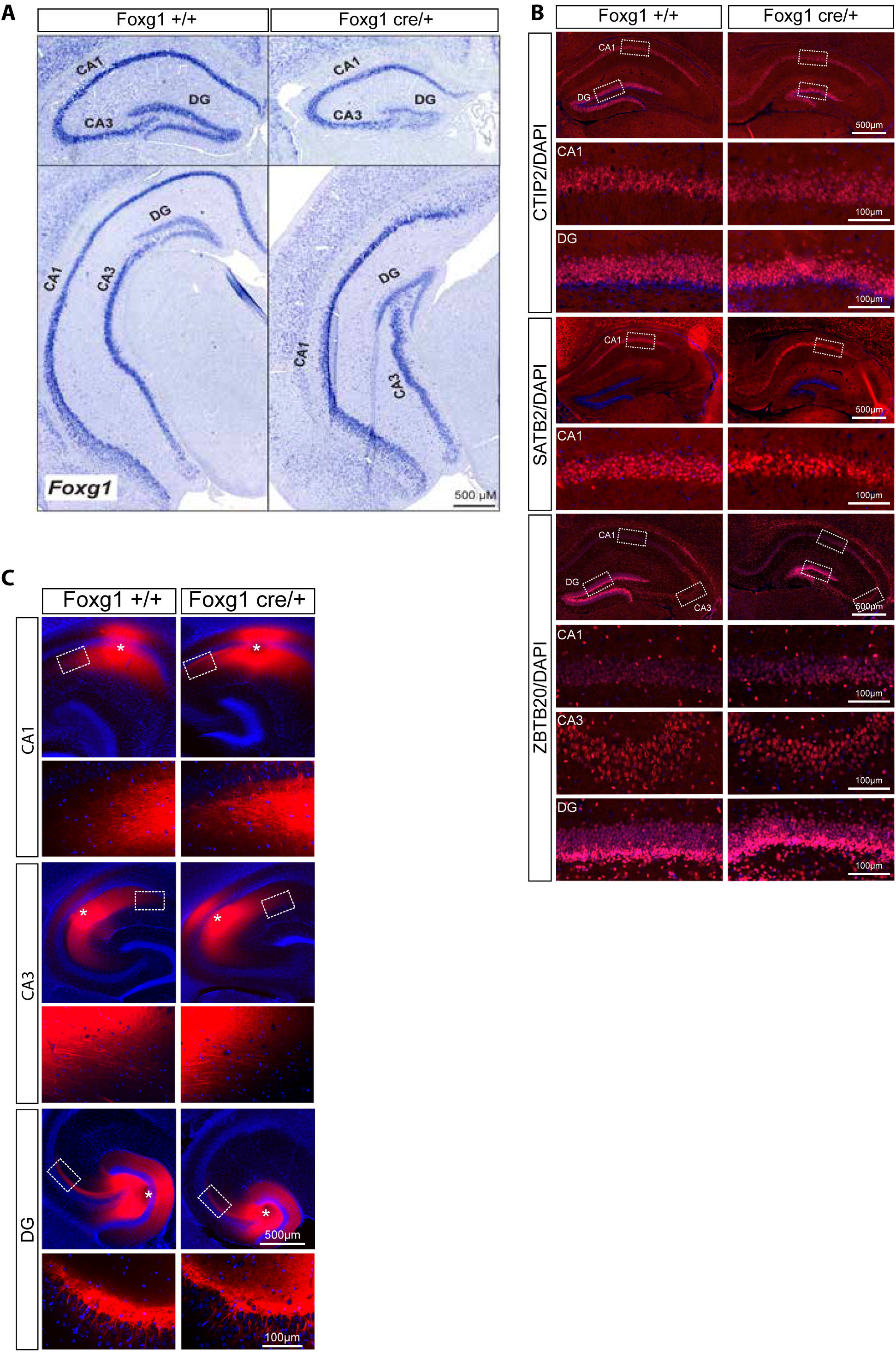
Histological characterization of the FOXG1-haploinsufficient adult mouse hippocampus. **(A)** Expression pattern of *Foxg1* in the 6-week old *Foxg1*^+/+^ (left, control) and *Foxg1*^cre/+^ (right) mouse hippocampus was detected by in situ hybridization. *Foxg1* was expressed in both the dentate gyrus (DG), and in the granule cells of the CA fields. *Foxg1*^cre/+^ mice had smaller hippocampus, although the CA and DG fields were preserved. n=1 **(B)** Immunostainings of 3-week old brains of *Foxg1*^cre/+^ and *Foxg1*^+/+^ for SATB2, CTIP2 and ZBTB20. The first row of images in each panel shows an overview of the hippocampus. The lower rows represent a magnification of the indicated regions in the overviews. n= 3 **(C)** Dil tracing in CA1, CA3, and DG in 7-week old mice shows that haploinsufficiency of FOXG1 did not affect intra- hippocampal connectivity. *Foxg1*^+/+^ n=3, *Foxg1^c^*^re/+^ n=5. CA: cornu ammonis, DG: dentate gyrus. Scale bars as indicated within the figures.

**Figure S2:**
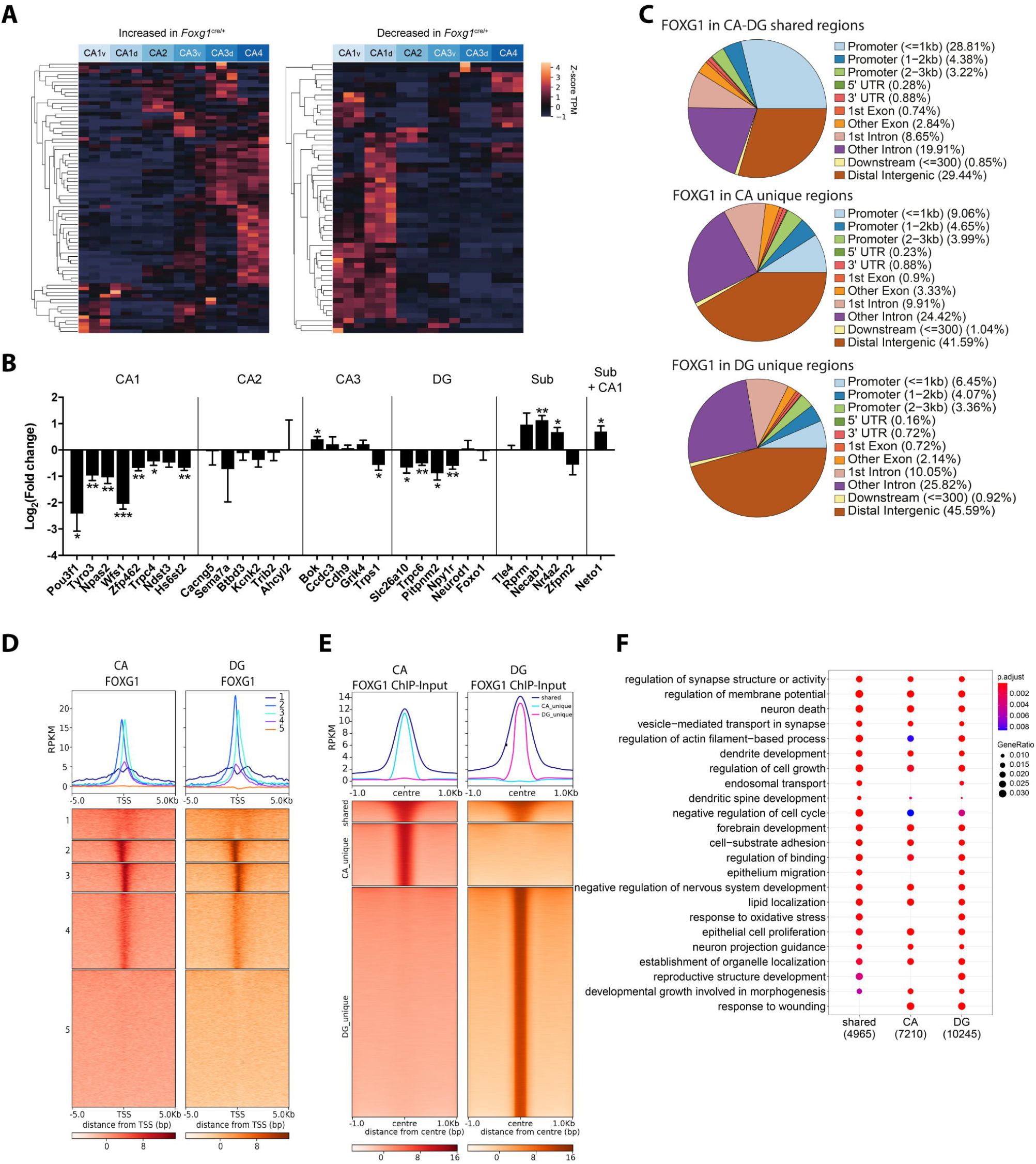
*Foxg1* haploinsufficiency affects gene expression in the CA fields and DG *in vivo*, but its cistrome is comparable *in vivo* and *in vitro*. (A) Genes that showed increased expression (left panel) in hippocampus of six-week old *Foxg1*^cre/+^ mice show relatively high expression in CA3/4. Genes that showed decreased expression (right panel) in hippocampus of six-week old *Foxg1*^cre/+^ mice show relatively high expression in CA1, as assessed with Hipposeq. **(B)** Expression analysis of marker genes for hippocampal subfields shows that CA1 and DG are most prominently affected after KD of FOXG1. qRTPCR analysis of selected marker genes for hippocampal subfields in hippocampus of six-week old *Foxg1*^cre/+^ mice compared to their respective wild type littermates shows that genes with high expression in CA1 or DG mostly decreased after KD of FOXG1. CA: cornu ammonis, DG: dentate gyrus, sub: subiculum. All qRTPCR data are represented as mean±SEM, n=3-5, *: p<0.05, **: p<0.01, ***p<0.001, unpaired Student’s t-test. **(C)** Genomic distribution of FOXG1 peaks in CA-DG shared, CA-unique, and DG-unique binding regions represented as pie charts. **(D)** K-means clustering (k=5) of FOXG1 enrichment in CA (red) and DG (orange) regions found 5 Kb up-/down-stream of TSS of protein coding genes. Data is normalized by sequencing depth and input control. The metaprofiles (top) show the average reads (RPKM) of each cluster. **(E)** Heatmap showing FOXG1 enrichment at binding sites clustered into shared, CA-unique, and DG-unique regions. Data normalization and metaprofiles (top) as in D. **(F)** GO-terms were compared among clusters derived from E. Scales of gene ratio and adjusted p-value are reported at the top right. Total number of genes per cluster are on the x-axis. Threshold for enrichment analysis was adjusted to p < 0.01.

**Figure S3:**
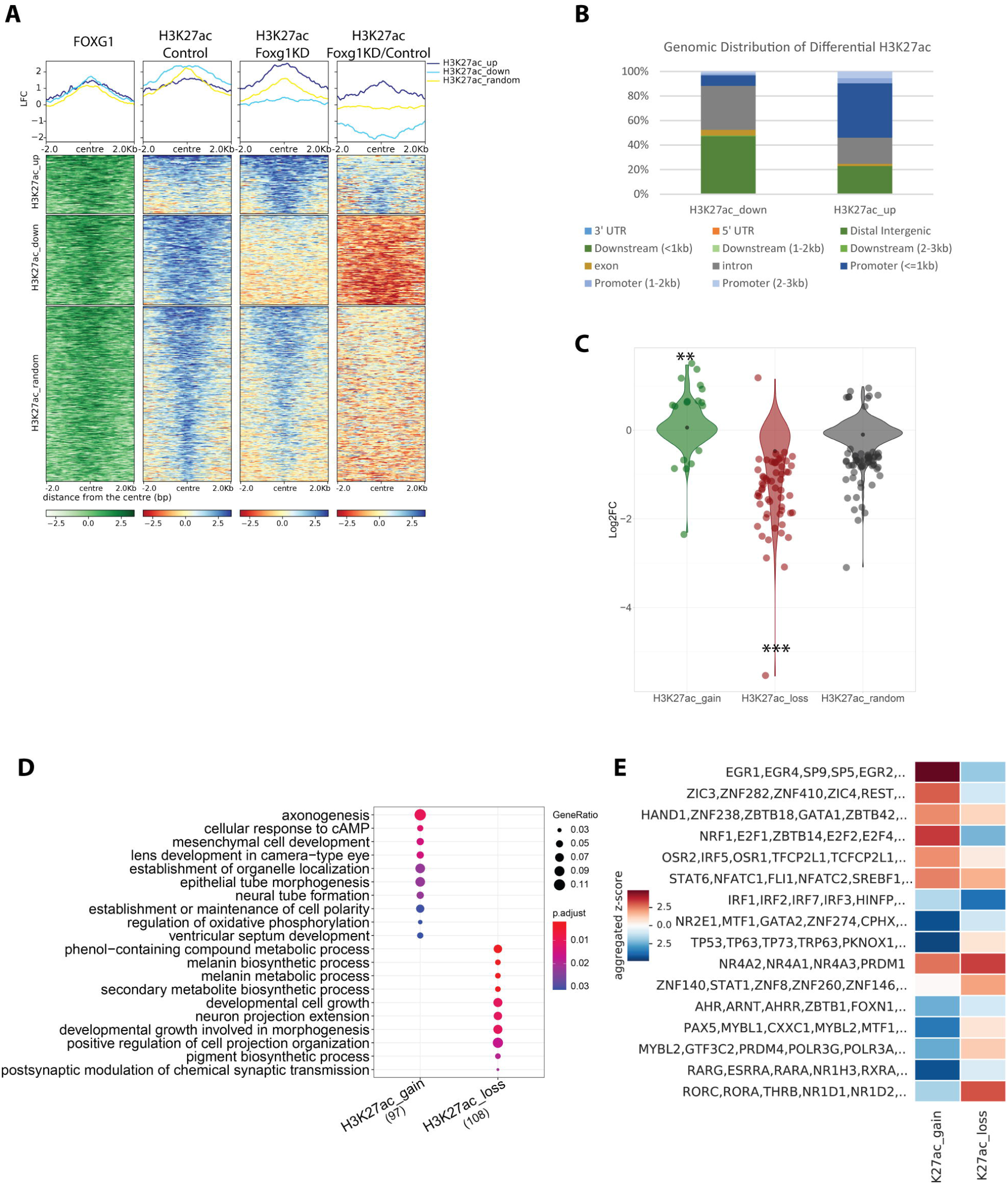
Gain and loss of H3K27ac upon reduced FOXG1 levels. **(A)** Heatmap of H3K27ac enrichment at regions retrieved from differential binding analysis of H3K27ac FOXG1 KD/Control (H3K27ac-up, -down, -random). Data is normalized by sequencing depth and input control as log_2_ (ChIP/Input) for H3K27ac control and H3K27ac FOXG1 KD data. The difference between FOXG1 KD and control conditions is calculated from RPKM normalized bigwig files as log_2_ (FOXG1 KD/Control). The metaprofiles (top) show the average log_2_FC (LFC) of each cluster. **(B)** Genomic distribution of regions gaining and losing H3K27ac enrichment displayed as a stacked bar graph. **(C)** Violin plot depicting the distribution of DEGs upon FOXG1 KD at H3K27ac-gain,–loss and -random clusters as shown in A. Y-axis corresponds to log_2_FC of gene expression, and x-axis shows the three clusters. The black dot marks the median of log_2_FC of DEGs in each cluster. Fisher’s exact test, *: p<0.05, **: p<0.01, ***: p<0.001. **(D)** Enriched GO-terms for the respective clusters as shown in A. Scales of gene ratios and adjusted p-value are at top-right corner, and total number of genes per cluster are on the x-axis. Threshold for enrichment analysis was adjusted to p < 0.01. **(E)** Heatmap showing transcription factor (TF)-binding differential motif analysis according to the clusters of H3K27ac enrichment as shown in A.

**Figure S4:**
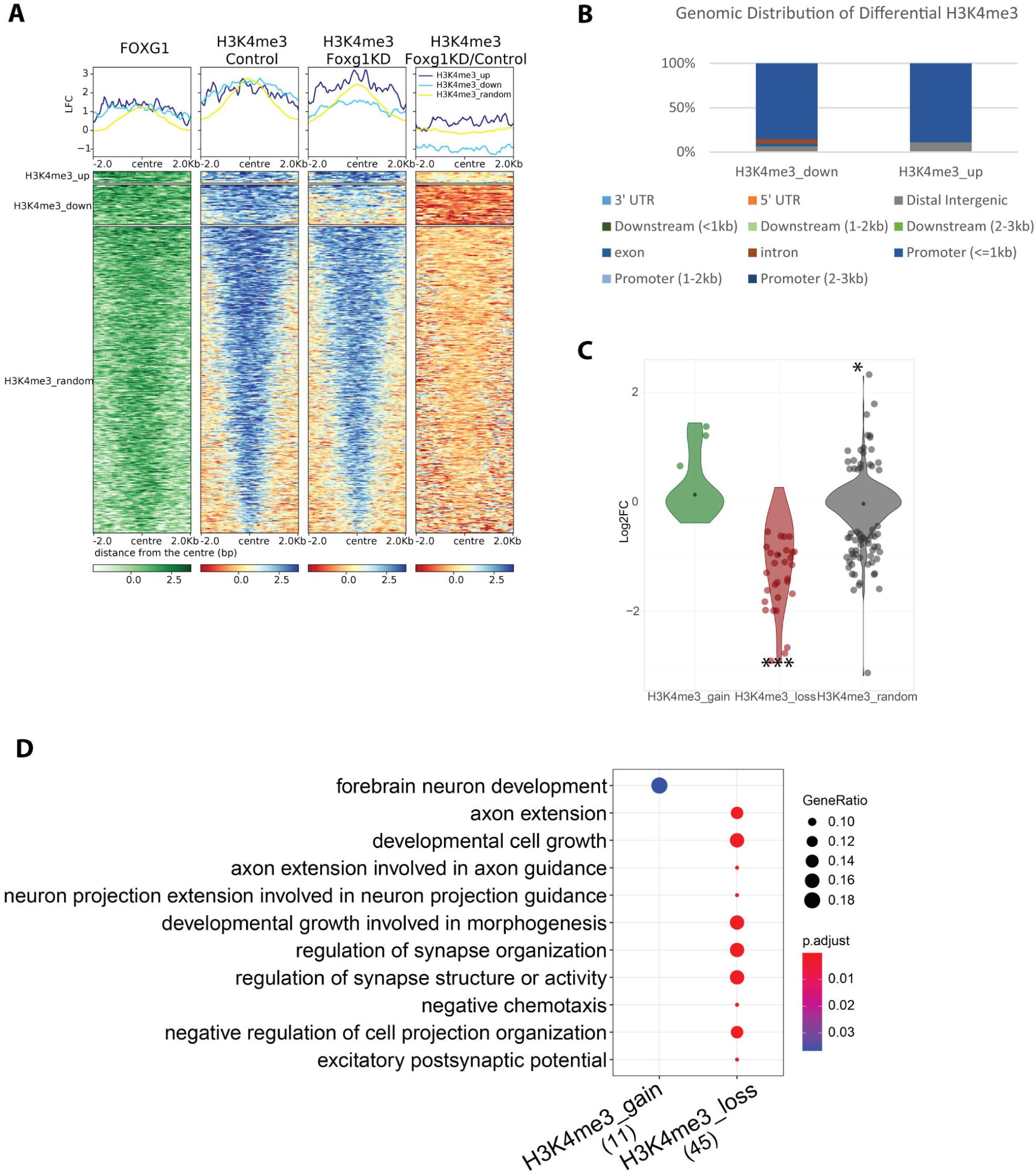
Gain and loss of H3K4me3 upon reduced FOXG1 levels. **(A)** Heatmap of H3K4me3 enrichment at regions retrieved from differential binding analysis of H3K4me3 FOXG1 KD/Control (H3K4me3-up, -down, -random). Data is normalized by sequencing depth and input control as log_2_ (ChIP/Input) for H3K4me3 control and H3K4me3 FOXG1 KD data. The difference between FOXG1 KD and control conditions is calculated from RPKM normalized bigwig files as log_2_ (FOXG1 KD/Control). The metaprofiles (top) show the average log_2_FC (LFC) of each cluster. **(B)** Genomic distribution of regions gaining and losing H3K4me3 enrichment displayed as a stacked bar graph. **(C)** Violin plot depicting the distribution of DEGs upon FOXG1 KD at H3K4me3-gain,–loss and -random clusters as shown in A. Y-axis corresponds to log_2_FC of gene expression, and x-axis shows the three clusters. The black dot marks the median of log_2_FC of DEGs in each cluster. Fisher’s exact test, *: p<0.05, **: p<0.01, ***: p<0.001. **(D)** Enriched GO-terms for the respective clusters as shown in A. Scales of gene ratios and adjusted p-value are at top-right corner, and total number of genes per cluster are on the x-axis. Threshold for enrichment analysis was adjusted to p < 0.01.

**Figure S5:**
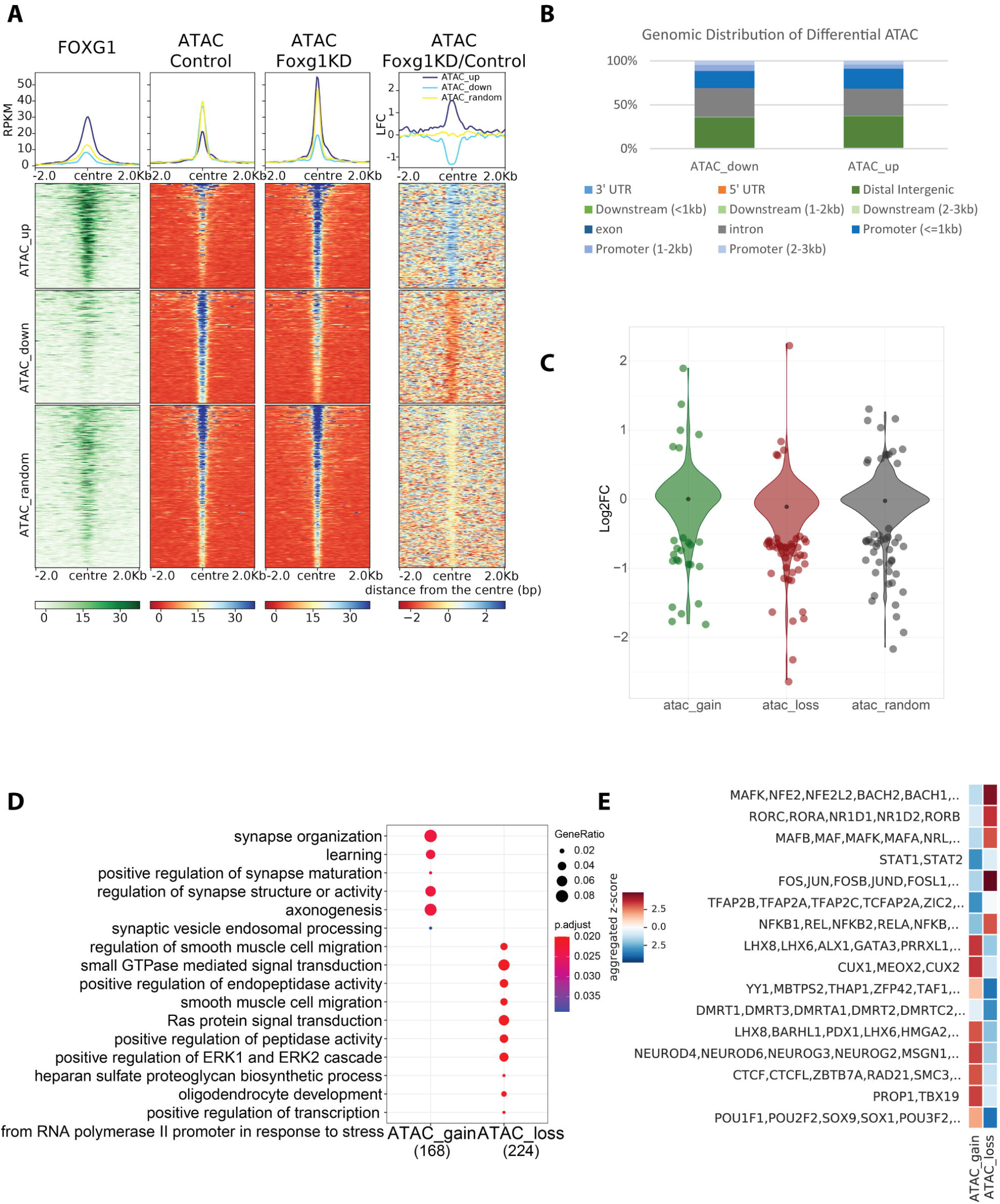
Gain and loss of accessibility upon reduced FOXG1 levels. **(A)** Heatmap of chromatin accessibility enrichment at regions retrieved from differential binding analysis of ATAC FOXG1 KD/Control (ATAC-up, -down, -random). Data is normalized by sequencing depth as RPKM for ATAC control and ATAC FOXG1 KD data. The difference between FOXG1 KD and control conditions is calculated from RPKM normalized bigwig files as log_2_(FOXG1 KD/Control). The metaprofiles (top) show the average log_2_FC (LFC) of each cluster. **(B)** Genomic distribution of regions gaining and losing accessibility displayed as a stacked bar graph. **(C)** Violin plot depicting the distribution of DEGs upon FOXG1 KD at ATAC-gain,–loss and -random clusters as shown in A. Y-axis corresponds to log_2_FC of gene expression, and x-axis shows the three clusters. The black dot marks the median of log_2_FC of DEGs in each cluster. Fisher’s exact test, *: p<0.05, **: p<0.01, ***: p<0.001. **(D)** Enriched GO-terms for the respective clusters as shown in A. Scales of gene ratios and adjusted p-value are at top-right corner, and total number of genes per cluster are on the x-axis. Threshold for enrichment analysis was adjusted to p < 0.01. **(E)** Heatmap showing transcription factor (TF)-binding differential motif analysis according to the clusters of chromatin accessibility as shown in A.

**Figure S6:**
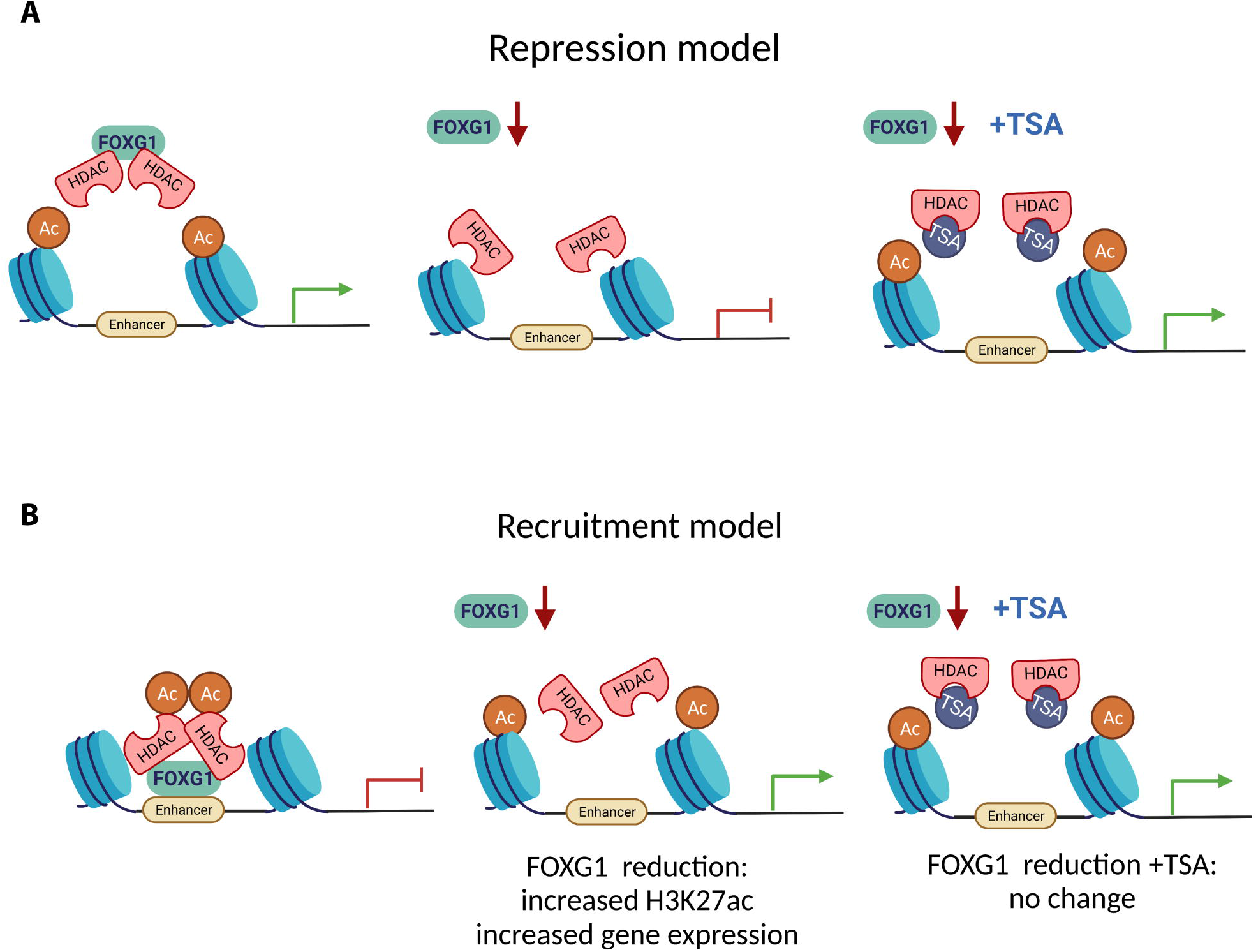
Repression and recruitment models of HDAC-FOXG1 interaction. Graphical summary of the repression **(A)** and recruitment **(B)** models**. (A)** The repression model is independent of FOXG1-binding to the chromatin and predicts that reduced levels of FOXG1 lead to reduced levels of H3K27ac and concomitant transcriptional decrease. Upon HDAC inhibition with TSA, we expect transcriptional increase of genes regulated through the repression model. **(B)** The recruitment model predicts that reduced levels of FOXG1 correlate with increased H3K27ac levels and concomitantly with increased transcription at FOXG1-binding. TSA treatment would not alter transcription upon reduced FOXG1 levels, as inhibition of the HDACs does not occur bound to the chromatin.

**Figure S7:**
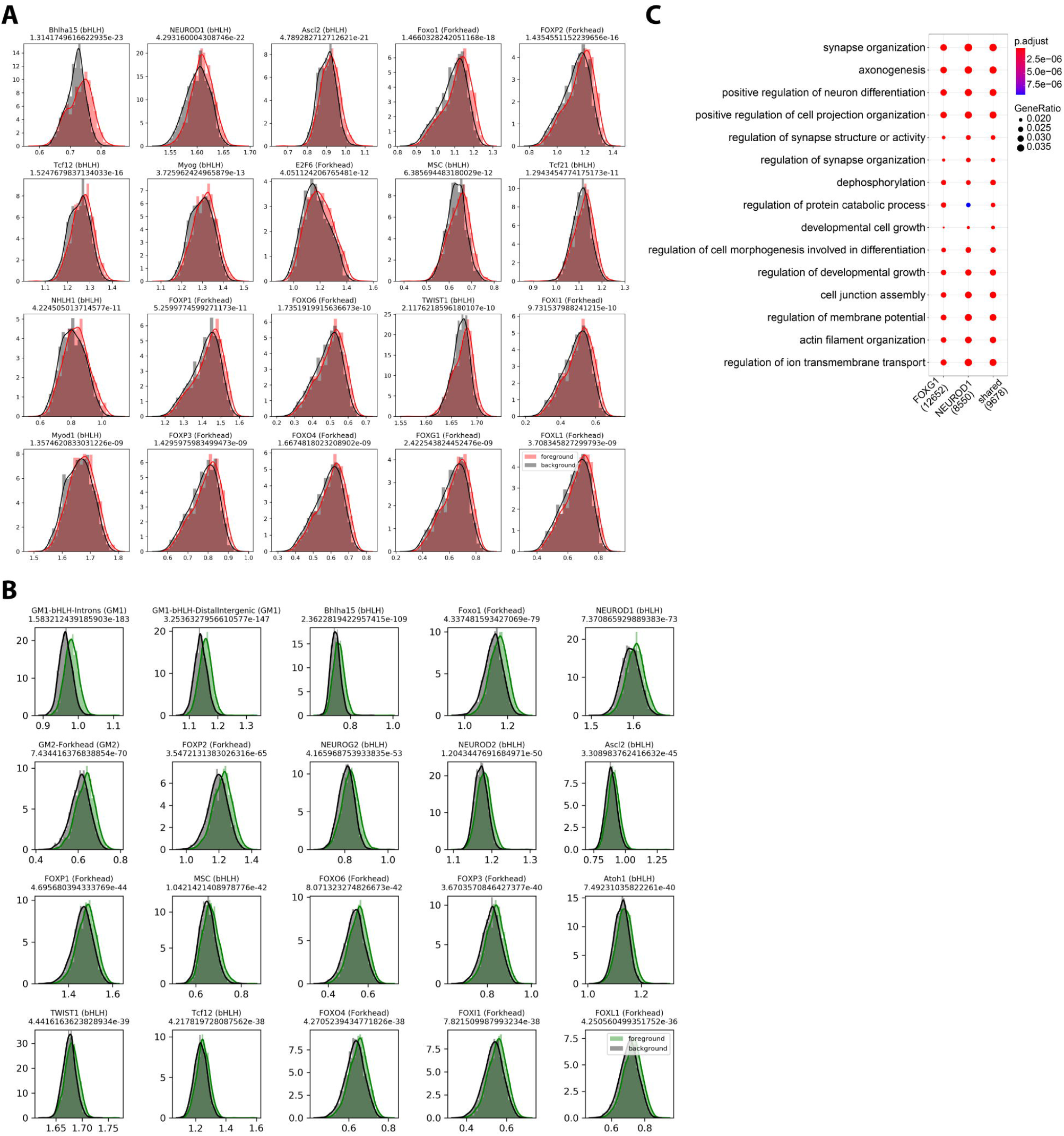
Motif affinity analysis of *in vitro* and *in vivo* FOXG1 peaks. Transcription factors that have the highest affinity for the sequences at +/− 200 bp flanking intronic and intergenic FOXG1 peak summits *in vivo* **(A)** and *in vitro* **(B)** as predicted by TRAP motif affinity analysis. **(C)** GO-term enrichment analysis of peaks shared between FOXG1 and NEUROD1, or unique to each TF. Scales of gene ratio and adjusted p-value reported at the top right. Total number of genes per cluster are on the x-axis. Threshold for enrichment analysis was adjusted to p < 0.01.

**Figure S8:**
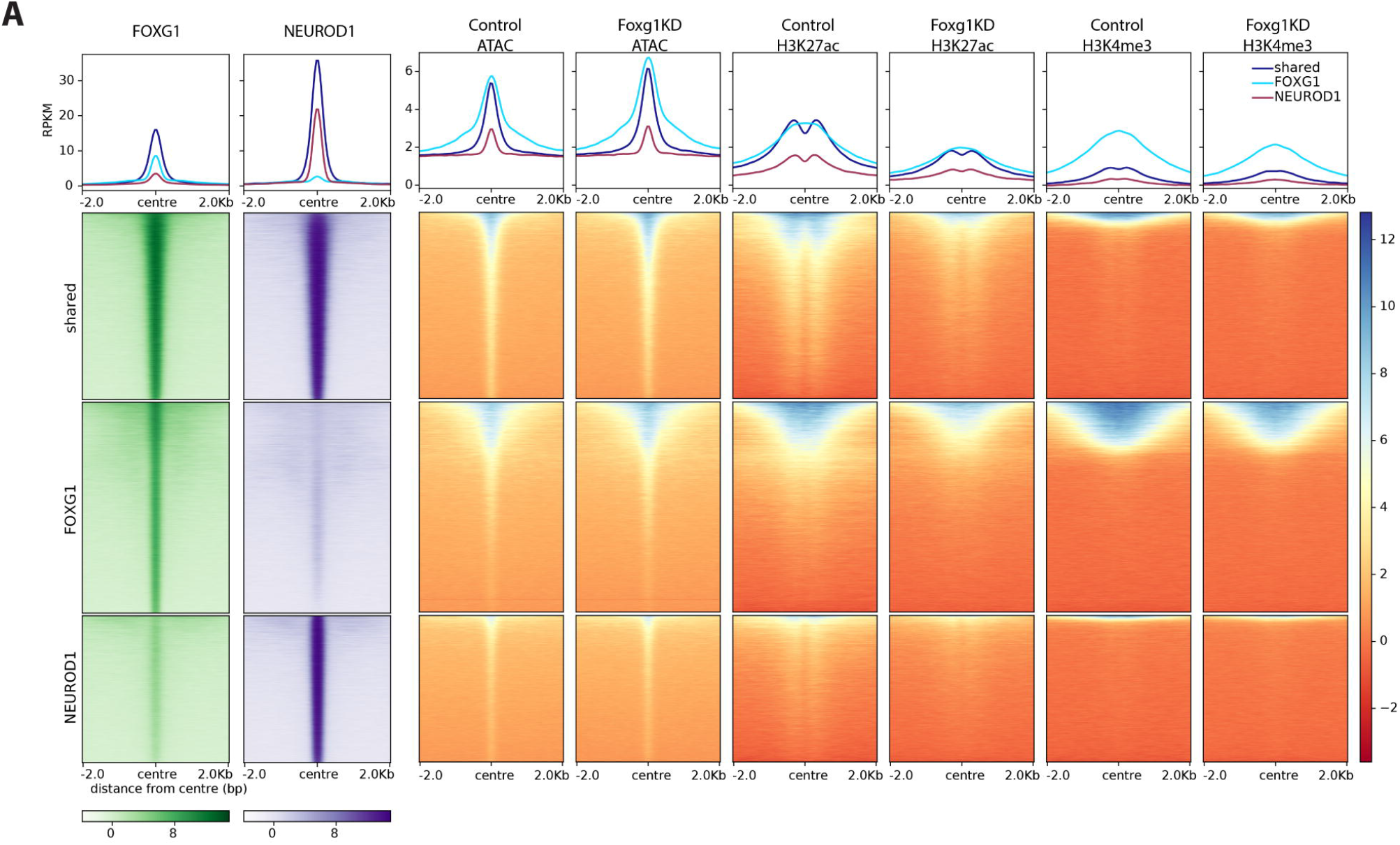
Epigenetic landscape at FOXG1 and NEUROD1 enriched regions. **(A)** Heatmap showing chromatin accessibility, H3K27ac, H3K4me3 enrichment at FOXG1 (green) and NEUROD1 (purple) binding sites clustered into shared and unique (FOXG1_unique, NEUROD1_unique) regions in control and FOXG1 KD conditions. Data normalization and metaprofiles (top) as in S5F.

## Declarations

### Ethics approval and consent to participate

not applicable

### Consent for publication

not applicable

### Availability of data and materials

The raw sequencing files were deposited to the NCBI Gene Expression Omnibus (GEO) and are available for download using the following accession number: GSExxxxx. All other data types and codes recreating the analyses from the data files can be found at http://www.github.com/ipekakol/FOXG1 as Jupyter notebooks or R markdown files. All other relevant data supporting the key findings of this study are available within the article and its Supplementary Information files or from the corresponding authors upon reasonable request.

### Author’s contribution

TV conceived and directed the project, and wrote the manuscript. IA, DoH, SH, AV, CH, TR performed experiments. CB executed the RELACS protocol and generated ChIP-seq data. DoH performed luciferase assays and library preparation for ATAC- and RNA-seq. TR performed sample and library preparation for RNA-seq. IA performed sample preparation for ChIP-seq. TM and AF facilitated the generation of high throughput data. IA performed all the analyses based on the high throughput datasets. TM coordinated analysis pipelines, and provided quality control and suggestions for data acquisition and analyses. TV, TM, IA, CH, DoH edited the manuscript. The authors read and approved the final manuscript.

## Acknowledgment

We thank F. Ferrari for his guidance in bioinformatics analyses. The authors thank B. Heimrich, G. Arumugam, E. Maier, and R. Vezzali for their technical help and input at different stages of the project. We also thank A. Izzo for input on immunoprecipitation experiments and critical comments on the manuscript. This work was supported by the Deutsche Forschungsgemeinschaft (DFG) through a grant to TV within the SPP1378 and through the DFG–fund 322977937/GRK2344. Further, the authors thank the Freiburg Galaxy Server team and all members of the Vogel group for discussion and support. We like to thank the team of the Sequencing Core at the MPI-IE for their support with data generation.

